# Safety, immunogenicity and protection provided by unadjuvanted and adjuvanted formulations of recombinant plant-derived virus-like particle vaccine candidate for COVID-19 in non-human primates

**DOI:** 10.1101/2021.05.15.444262

**Authors:** Stéphane Pillet, Prabhu S. Arunachalam, Guadalupe Andreani, Nadia Golden, Jane Fontenot, Pyone Aye, Katharina Röltgen, Gabrielle Lehmick, Charlotte Dubé, Philipe Gobeil, Sonia Trépanier, Nathalie Charland, Marc-André D’Aoust, Kasi Russell-Lodrigue, Robert V. Blair, Scott Boyd, Rudolph B. Bohm, Jay Rappaport, François Villinger, Brian J. Ward, Bali Pulendran, Nathalie Landry

**Affiliations:** Medicago Inc., Québec, QC, Canada; Institute for Immunity, Transplantation and Infection, Stanford University School of Medicine, Stanford University, Stanford, CA, USA; Tulane National Primate Research Center, Covington, LA, USA; New Iberia Research Center, University of Louisiana at Lafayette, New Iberia, LA, USA; Department of Pathology, Stanford University School of Medicine, Stanford University, Stanford, CA, USA; Department of Microbiology and Immunology, Tulane University School of Medicine, New Orleans, LA, USA; Research Institute of the McGill University Health Centre, Montreal, QC, Canada; Department of Microbiology and Immunology, Stanford University School of Medicine, Stanford University, Stanford, CA, USA; Institute for Immunity, Transplantation & Infection, Stanford University School of Medicine, Stanford University, Stanford, CA, USA

**Keywords:** SARS-CoV-2, COVID-19 vaccine, Virus-like particles, safety, immunogenicity, protection, non-human primates

## Abstract

Although antivirals are important tools to control the SARS-CoV-2 infection, effective vaccines are essential to control the current pandemic. Plant-derived virus-like particle (VLP) vaccine candidates have previously demonstrated immunogenicity and efficacy against influenza. Here we report the immunogenicity and protection induced in macaques by intramuscular injections of VLP bearing SARS-CoV-2 spike protein (CoVLP) vaccine candidate formulated with or without Adjuvant System 03 (AS03) or cytosine phosphoguanine (CpG) 1018. Although a single dose of unadjuvanted CoVLP vaccine candidate stimulated humoral and cell-mediated immune responses, booster immunization (at 28 days after prime) and adjuvants significantly improved both responses with a higher immunogenicity and protection provided by AS03 adjuvanted CoVLP. Fifteen microgram CoVLP adjuvanted with AS03 induced a balanced IL-2 driven response along with IL-4 expression in CD4 T cells and mobilization of CD4 follicular helper cells (Tfh). Animals were challenged by multiple routes (i.e. intratracheal, intranasal and ocular) with a total viral dose of 10^6^ plaque forming units of SARS-CoV-2. Lower viral replication in nasal swabs and broncho-alveolar lavage (BAL) as well as fewer SARS-CoV-2 infected cells and immune cell infiltrates in the lungs concomitant with reduced levels of pro-inflammatory cytokines and chemotactic factors in BAL were observed in the animals immunized with CoVLP adjuvanted with AS03. No clinical, pathologic or virologic evidences of vaccine associated enhanced disease (VAED) were observed in vaccinated animals. CoVLP adjuvanted with AS03 was therefore selected for vaccine development and clinical trials.

## Introduction

In December 2019, a series of severe atypical respiratory disease cases occurred in Wuhan China, and a novel coronavirus named Severe Acute Respiratory Syndrome Coronavirus 2 (SARS-CoV-2) was rapidly identified as the causative agent of coronavirus disease 2019 (COVID-19). SARS-CoV-2 virions consist of a helical nucleocapsid, formed by association of nucleocapsid (N) phosphoproteins with viral genomic RNA that is surrounded by a lipid bilayer, into which three structural proteins are inserted: the spike (S), the membrane (M) and the envelope (E) proteins. The new coronavirus rapidly spread around the globe resulting in the World Health Organization’s declaration of a pandemic on March 11, 2020. As of May 5, 2021, SARS-CoV-2 has caused over 155 million infections and more than 3.2 million deaths worldwide (Johns Hopkins University of Medicine Coronavirus Resource Center. COVID-19 dashboard). Although therapeutics including monoclonal antibodies and antivirals have shown some therapeutic efficacy in limiting the burden of COVID-19, effective vaccines are essential to control the pandemic. As of April 20, 13 vaccines were approved by different Health Agencies worldwide and over 10 vaccine candidates are in phase 2/3 or phase 3, including the plant-based vaccine candidate developed by Medicago.

The S glycoprotein contains a receptor binding domain (RBD) that binds strongly to human angiotensin-converting enzyme 2 (ACE2) receptors playing a major role in viral attachment, fusion and entry into host cells^1,2^. Neutralizing antibodies (NAb) directed against the S protein were known to provide protection from other highly pathogenic coronaviruses (e.g.: SARS-1, MERS) and similar protective efficacy was rapidly demonstrated with anti-S antibodies in SARS-CoV-2 infection^3,4^. Consequently, the S protein rapidly emerged as a prime target for the development of COVID-19 vaccines.

Medicago’s platform technology is based on transient expression of recombinant proteins in the non-transgenic plant *Nicotiana benthamiana. Agrobacterium tumefaciens* is used as a transfer vector to move targeted DNA constructs into the plant cells. The newly introduced DNA then directs the expression of the desired recombinant proteins^5^. Plant-derived coronavirus-like particles (CoVLP) are produced from the expression of a modified full-length S protein. In plant cells, the newly synthesized S proteins trimerize and then move to lipid rafts of the plasma membrane where they spontaneously assemble into virus-like particles (VLPs) that ‘bud’ off the surface of the plant cell. The S proteins in CoVLP are in a stabilized, prefusion conformation that resembles the native structure seen on SARS-CoV-2 virions. The prefusion form of S protein is preferred as a vaccine antigen since it contains several epitopes in the RBD that are primary targets for neutralising antibodies^6^. Moreover, previous study on the MERS S protein revealed that the prefusion state also resulted in a more potent immunogen with dose-sparing properties compared to protein made with the original wild-type sequence^7^. In a pandemic situation, large numbers of vaccine doses are required to protect the maximum number of people worldwide. Dose-sparing adjuvants are often used to maximize the number of vaccine doses available, i.e. to reduce the amount of antigen needed to elicit a robust and persistent immune response^8^. Two adjuvants with demonstrated potential for dose-sparing were included in the current study: the cytidine-phospho-guanosine (CpG)-containing immunostimulatory oligodeoxynucleotide sequence 1018 (CpG 1018: Dynavax) and the α-tocopherol-containing oil-in-water emulsion Adjuvant System 03 (AS03: GSK).

Synthetic oligodeoxynucleotides like CpG 1018 are toll-like receptor 9 (TLR9) agonists mimicking the activity of naturally occurring CpG motifs found in bacterial DNA. CpG are potent vaccine adjuvants that enhance immune responses in general and tend to promote Th1-type responses in particular. B cells and plasmacytoid dendritic cells are the main human immune cells that express TLR9 and activation of these cells by CpG DNA initiates an immunostimulatory cascade that culminates in the indirect maturation, differentiation and proliferation of natural killer (NK) cells, T cells and monocytes/macrophages that contribute to a Th1-biased immune response^9,10^. CpG 1018 is the adjuvant used in the licensed hepatitis B vaccine HEPLISAV-B.

The Adjuvanted System AS03 initiates a transient innate immune response at the injection site and draining lymph node in animal models^11^ and in human peripheral blood^12–14^. That innate immune response potentiates and shapes the adaptive immune response to the vaccine antigen including both antibody and T-cell responses, resulting in increased response magnitude, breadth, durability and antibody avidity^15–17^. AS03 has been used in the licensed pandemic A/H1N1pdm09 influenza vaccines Arepanrix H1N1 (in Canada), Q Pan H5N1 (in the USA) and Pandemrix (in Europe), of which 90 million doses have been administered worldwide, as well as in other licensed (Q Pan H5N1 in the USA) or vaccine candidates^18^. The present study assessed the safety, immunogenicity, and protective efficacy of two doses, 28-days apart, of a plant-produced virus-like-particle vaccine candidate for COVID-19 in rhesus macaques.

## Material and Methods

### Animal model

Male Indian rhesus macaques from 3.5 to 8 years old were sourced from the breeding colonies of the New Iberia Research Center (NIRC) of the University of Louisiana et Lafayette and distributed evenly among the treatment groups based on their age and weight to assure comparable distribution across treatments. All animals were negative for simian immunodeficiency virus (SIV), simian T cell leukemia virus, simian retrovirus and Herpes B virus. Animals were housed and maintained as per National Institutes of Health (NIH) guidelines at NIRC of the University of Louisiana at Lafayette in accordance with the rules and regulations of the Committee on the Care and Use of Laboratory Animal Resources. The entire study (protocol 2020-8721-007) was reviewed and approved by the University of Louisiana at Lafayette Institutional Animal Care and Use Committee (IACUC). For the challenge, animals were transferred to the Regional Biosafety Level 3 (BSL3) facility at the Tulane National Primate Research Center, where the study was reviewed and approved by the Tulane University IACUC (Protocol P0450). All animals were cared for in accordance with the Institute for Laboratory Animal Research Guide for the Care and Use of Laboratory Animals 8th Edition. The Tulane University Institutional Biosafety Committee approved the procedures for sample handling, inactivation, and removal from BSL3 containment.

### Vaccine and adjuvants

The full-length S glycoprotein of SARS-CoV-2, strain hCoV-19/USA/CA2/2020, corresponding in sequence to nucleotides 21563 to 25384 from EPI_ISL_406036 in GISAID database (https://www.gisaid.org/) was transiently expressed in *Nicotiana benthamiana* plants using a similar strategy previously described to produce influenza vaccine candidate^5^. The S protein was modified with R667G, R668S and R670S substitutions at the S1/S2 cleavage site to increase stability, and K971P and V972P substitutions to stabilize the protein in the prefusion conformation^19^. The signal peptide was replaced with a plant gene signal peptide and the transmembrane domain (TM) and cytoplasmic tail (CT) of S protein were also replaced with TM/CT from Influenza hemagglutinin H5 from A/Indonesia/5/2005 to increase VLP assembly and budding. The self-assembled VLPs bearing S protein trimers were isolated from the plant matrix and subsequently purified using a process similar to that described for the influenza vaccine candidates^5^. AS03 (provided by GSK, Rixensart, Belgium) is an o/w emulsion containing 11.86 mg DL-α-tocopherol, 10.69 mg squalene and 4.86 mg polysorbate 80 per 250 μL, which corresponds to one human dose. CpG 1018, a synthetic 22-based phosphorothioate oligodeoxynucleotide containing a cytidine-phospho-guanosine (CpG) immunostimulatory sequence in sterile Tris buffered saline solution, was provided by Dynavax (Emeryville, CA, USA).

### Study design

Male Indian rhesus macaques were immunized intramuscularly (IM) with one (at day 0, D0) or two (D0 and day 28, D28) doses of 15 μg of CoVLP (formulated with or without CpG 1018 or AS03) or control phosphate buffered solution. Approximately one week before challenge, the animals were transferred to the Tulane National Primate Research Center (TNPRC) for viral challenge. Animals were infected (detailed below) with SARS-CoV-2 USA-WA1/2020 on D28 (one dose groups) or D57 (two dose groups) and euthanized 6 or 20 days after challenge (Figure 1).

**Figure 1:**
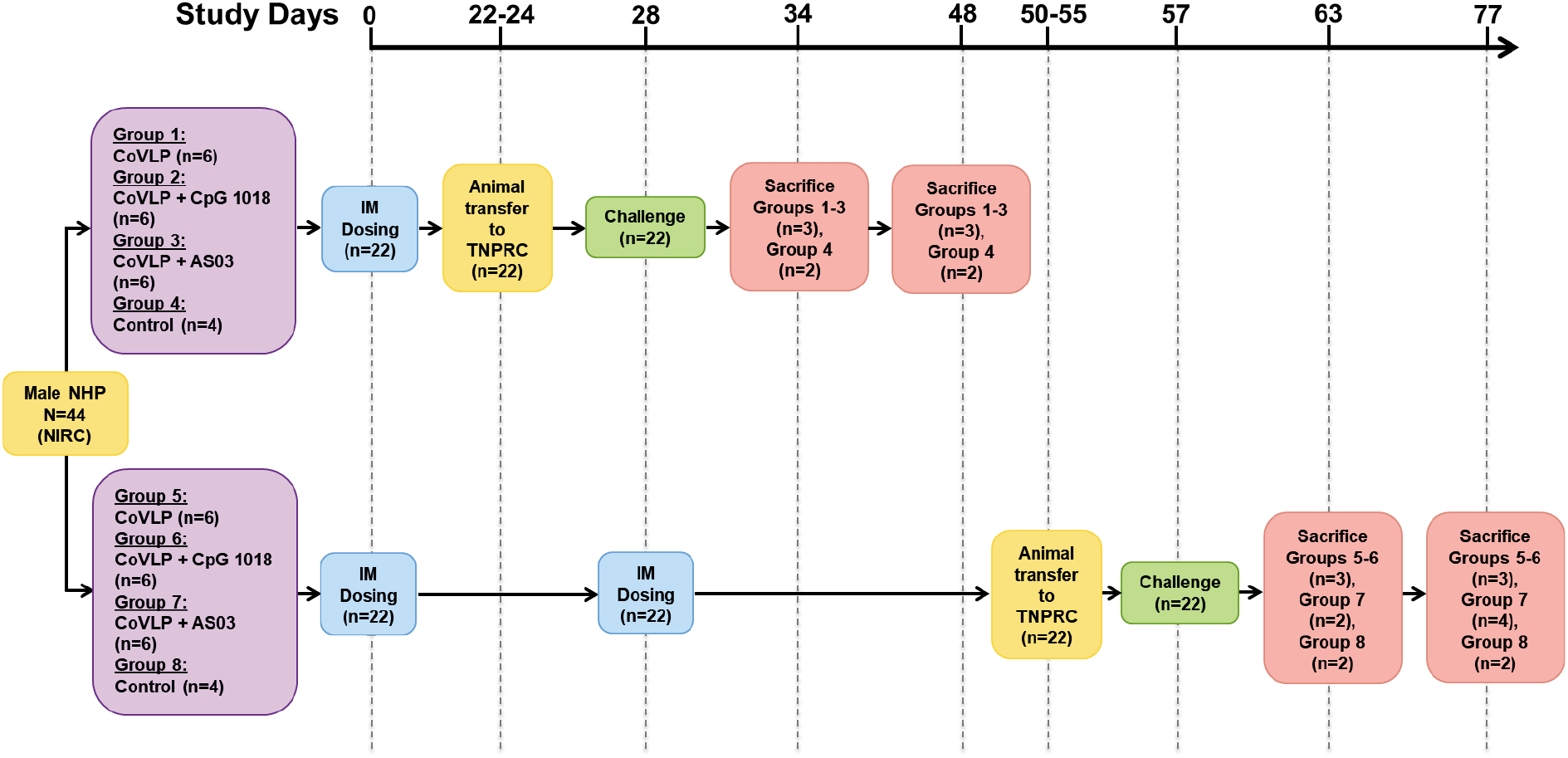
Experimental design

Treatment groups are detailed in Suppl. Table 1. For logistic reasons, the study was carried out in two cohorts of four groups each (i.e. the One-dose cohort included Groups 1-4 and the Two-dose Cohort included Groups 5-8): animals in Groups 4 and 8 received placebo (Control); Groups 1 and 5 received CoVLP (15 μg in 0.5 mL/dose) without an adjuvant (CoVLP); Groups 2 and 6 received CoVLP (15 μg/dose) formulated with CpG 1018 (3 mg/dose) for a final volume of 1 mL (CoVLP+CpG); Groups 3 and 7 received CoVLP (15 μg/dose) formulated with AS03 (1:1 ratio v/v, CoVLP+AS03) for a final volume of 0.5 mL. Adjuvanted vaccines were mixed immediately prior to dosing and all vaccines were administered IM in the deltoid muscle.

### Pre-challenge safety evaluation

#### Clinical Observations

Clinical observations, including general behavior, visible signs of disease (reduced appetite; hunched posture; pale appearance; dehydration) were performed daily before challenge.

#### Body Temperature and Weight

Body temperature and weight were measured for all animals at each scheduled anesthetic event, i.e. on D0 (before immunization), D1, D3 and D7 after the first immunization for all groups and on D28 (before second immunization), D29, D32 and D35 for groups 5 to 8. Body temperature and weight changes are presented as relative to the baseline before each immunization.

#### Injection Site Observations

The injection sites were monitored for 7 days after each immunization using standard measures including upper arm circumference, lymph node enlargement (size, mobility, number), warmth, erythema, local lesions (ulcers, abscess) or other abnormalities. Each of these parameters was graded throughout the study by the same experienced investigator blind to group assignment (0 = absent, 1 = mild, 2 = moderate, 3 = severe). Upper arm circumference and lymph node enlargement were manually assessed at scheduled anesthetic events, up to 7 days after injection (D0, D1, D3 and D7 for all groups and D28 (before dosing), D29, D32 and D35 for groups 5 to 8).

#### Blood Biochemistry and Hematology

Routine hematologic (e.g. hematocrit, hemoglobin level, and white blood cell count) and biochemical analyses (e.g. blood urea nitrogen, serum creatinine, total protein, albumin, aspartate aminotransferase, gamma-glutamyl transpeptidase, and C-reactive protein [CRP]) were performed on whole blood samples from all animals. Blood samples were collected before immunization (baseline) and on D1, D3 and D7 for the One-dose cohort; the day of the second immunization (before dosing), D29, D32 and D35 for the Two-dose cohort.

### Humoral Immune Response

#### NAb Titers

Serum samples from all animals were collected at baseline, D21 and D49. NAb titers were measured by laboratory staff blind to group allocation at Battelle Biomedical Research Center (OH, USA) using a SARS-CoV-2 microneutralization (MN) assay with an *in situ* ELISA readout against SARS-CoV-2 NP. Briefly, serum samples were incubated with SARS-CoV-2 WA 1/2020 isolate at 37.0±2°C and 5.0±2% CO_2_ for 60±5 minutes. The serum/virus mixture was then transferred to previously seeded VERO E6 cell monolayers and incubated for 40 to 46 hours after which the inoculum was removed and plates were rinsed with Hank’s Balanced Salt Solution (HBSS) prior to fixation with 80% acetone. Following fixation, an *in situ* ELISA was performed for the detection of SARS-CoV-2 NP antigen using an anti-SARS-COV-2, MERS, SARS-CoV NP primary antibody (EastCoast Bio; ME, USA) and a Goat Anti-Mouse IgG conjugate antibody (Fitzgerald; MA USA). Samples were analyzed in duplicate and results are reported as the median of two independent analyses. The final value for each sample is expressed as MN50. Nonresponders (MN50 value <20), if any, were assigned a value of 10.

#### IgG, IgM and IgA Antibody Titers

Levels of vaccine-induced anti-Receptor Binding Domain (RBD) specific immunoglobulin (Ig)G, IgA and IgM in serum were measured in using an ELISA method similar to that described by Roltgen and co-authors^20^. RBD-specific IgG, IgA and IgM were measured at baseline, D14, D21 (all groups); and at D28, D42 and D49 (Two-dose cohort). Briefly, 96-well plates were coated with SARS-CoV-2 RBD protein. Following washing and blocking steps, wells were incubated with NHP serum samples at a dilution of 1/100. The dilution of 1/100 was determined in method optimization covering a 1/50 to 1/25600 dilution range. Four negative sera were also included in the optimization process. After washing to remove unbound antibodies, specific RBD-bound IgG, IgA or IgM antibodies were detected by horseradish peroxidase-conjugated goat anti-monkey/human IgG (γ-chain specific, catalog no. 62-8420, Thermo Fisher), IgM (μ-chain specific, catalog no. A6907, Sigma), or IgA (α-chain specific, catalog no. P0216, Agilent) secondary antibodies. After washes, 3,3’,5,5’-Tetramethylbenzidine (TMB) substrate solution was added, and the reaction was stopped using sulfuric acid. The optical density (OD) at 450 nanometers was measured with an EMax Plus microplate reader (Molecular Devices, San Jose, CA); values for blank wells were subtracted from values obtained for sera samples. All samples were run in duplicate.

### Cell-mediated immune response

Antigen-specific T cell responses were measured by intracellular cytokine stain (ICS) assay at baseline, D7 and D21 after the first dose; and D35 and D49 after the second dose. Frozen PBMCs were thawed, counted and resuspended at a density of 1 million live cells/mL in complete RPMI (RPMI supplemented with 10% FBS and antibiotics). The cells were rested overnight at 37°C in CO_2_ incubator. Next morning, the cells were counted again, resuspended at a density of 15 million/mL in complete RPMI and 100 μL of cell suspension containing 1.5 million cells was added to each well of a 96-well round-bottomed tissue culture plate and stimulated *ex vivo* with a peptide pool consisting of 15mer peptides overlapping by 11 amino acids spanning the S protein (GenScript, Piscataway, NJ), at a concentration of 1.2 μg/mL of each peptide in the presence of 1 μg/mL anti-CD28. The samples were incubated at 37°C in a CO_2_ incubators for 2h before addition of 10 μg/mL Brefeldin-A. The cells were incubated for an additional 4h. The cells were washed with PBS and stained with Zombie UV fixable viability dye (BioLegend). The cells were washed with PBS containing 5% FCS, before the addition of surface antibody cocktail (Suppl. Table 2). The cells were stained for 20 min at 4°C in 100 μL volume. Subsequently, the cells were washed, fixed and permeabilized with cytofix/cytoperm buffer (BD Biosciences) for 20 minutes. The permeabilized cells were stained with ICS antibody cocktail (Suppl. Table 2) for 20 min at room temperature in 1X-perm/wash buffer (BD Biosciences). Cells were then washed twice with perm/wash buffer (BD Biosciences) and once with staining buffer before analysis using BD Symphony Flow Cytometer. All flow cytometry data were analyzed using Flowjo software v10 (TreeStar Inc.).

### Challenge

Animals were infected under anesthesia with viral isolate SARS-CoV-2 USA-WA1/2020 (originally obtained from a patient in Seattle, USA, kindly provided by University of Texas, Medical Branch, Galveston, Texas) as previously described^21,22^. The virus stock was generated by expansion of a seed stock on Vero E6 cells and titrated by plaque assay on Vero E6 cells. It was deep sequenced and found to contain no polymorphisms at greater than 5% of reads relative to the original patient isolate. The furin cleavage site, a site with frequent culture adaptation in Vero E6 cells, harbored no polymorphisms at greater than 1% of sequence reads in this stock. Animal were challenged by multi-route administration (i.e. 500 μL intratracheal, intranasal (approximately 250 ul on each nares) and conjunctival (10 μL)) with a total viral dose of 10^6^ plaque forming units (PFU).

### Protection and Post-challenge safety evaluation

#### Clinical Observations and chest radiographs

Clinical observations, including state of responsiveness, presence of discharge, skin condition, respiratory condition, food consumption and fecal conditions were performed daily after challenge until termination by laboratory staff blind to group allocation. Animals were individually scored for the severity of these parameters and clinical scores were assigned according to the chart provided in supplementary material (Appendix). Thoracic radiographs were obtained 6, 13 and 20 days post-challenge.

#### Body Temperature, Weight, Respiratory Rate and Oxygen Saturation

Body temperature, weight, respiratory rate and levels of oxygen saturation (SpO2) were measured for all animals on the challenge day (before virus administration) and on days 6, 13 and 20 post-challenge for all animals. Body temperature, weight and SpO2 changes are presented as relative to challenge day (before infection).

#### Blood Biochemistry and Hematology

Routine hematologic (e.g. hematocrit, hemoglobin level, and white blood cell count) and biochemical analyses (e.g. blood urea nitrogen, serum creatinine, total protein, albumin, aspartate aminotransferase, gamma-glutamyl transpeptidase, and CRP) were performed on whole blood samples from all animals 6, 13 and 20 days post-challenge.

#### Gross Pathology Examination

Tissue (lungs, liver, kidneys, spleen, thymus and heart) samples for routine gross pathology examination were collected from all animals, i.e. 3 animals per group on two occasions, 6 and 20 days post viral infection, with the exception of the CoVLP+AS03 group that received 2 immunizations (group 7) from which 2 animals were euthanized 6 days after infection and 4 animals 20 days after infection. Animals were chosen randomly.

#### Viral replication

Viral replication was assessed by measuring subgenomic RNA (sgRNA) levels of E and/or N gene of SARS-CoV-2 by real time polymerase chain reaction (PCR) in broncho-alveolar lavage (BAL) as well as in nasal, oropharyngeal and anal swabs 6, 13 and 20 days after viral infection in all animals. Samples were collected in 200 μL of DNA/RNA Shield 1x (Zymo Research, Irvine, CA) and extracted for Viral RNA (vRNA) using the Quick-RNA Viral kit (Zymo Research). The Viral RNA Buffer was dispensed directly to the swab in the DNA/RNA Shield. A modification to the manufacturers’ protocol was using which involved insertion of the swab directly into the spin column for centrifugation allowing all of the collected material to cross the spin column membrane. The vRNA was the eluted (45 μL) from which 5 μL was added in a 0.1 mL fast 96-well optical microtiter plate format (Thermo Fisher, CA) for a 20 μL RT-qPCR reaction. The RT-qPCR reaction used TaqPath 1-Step Multiplex Master Mix (Thermo Fisher) along with 2019-nCoV RUO Kit (IDTDNA, Coralville, IA) a premix of forward and reverse primers and a FAM labeled probe targeting the N1 amplicon of N gene or a leader sequence on the *E* gene of SARS2-nCoV19 (accession MN908947). The reaction master mix was added to the microtiter plates using an X-stream repeating pipette (Eppendorf, Hauppauge, NY) which were then covered with optical film (Thermo Fisher), vortexed, and pulse centrifuged. The RT-qPCR reaction was subjected to RT-qPCR a program of UNG incubation at 25°C for 2 minutes, RT incubation at 50°C for 15 minutes, and an enzyme activation at 95°C for 2 minutes followed by 40 cycles of a denaturing step at 95°C for 3 seconds and annealing at 60°C for 30 seconds. Fluorescence signals were detected with an Applied Biosystems QuantStudio 6 Sequence Detector. Data were captured and analyzed with Sequence Detector Software v1.3 (Applied Biosystems, Foster City, CA). The baseline was adjusted from 15 to 10 Cq based on the 10^8^ copy number of the calibration curve intersecting with the threshold line at 12 (± 0.5) cycles. Viral copy numbers were determined by plotting Cq values obtained from unknown (i.e. test) samples against Serial tenfold dilutions of an exogenous calibration curves ranging from 10^1^ to 10^8^ log copy numbers generated from known amounts of *in vitro*-transcribed ssRNA for interpolation to determine the number of virus copies per mL of initial specimen sample or test dilution, as applicable. The limit of detection of the assay was 10 copies per reaction volume. To control for assay variation, validation of results, IPC quantification consistency, and reagent stability, replicates of a positive control sample also were analyzed in parallel with every set of test samples. Negative control plasma samples were included for processing with every set of test samples to monitor cross-contamination. A non-template control (NTC) was included in all runs to ensure that there was no cross-contamination between reactions.

#### Chemokines and Cytokines in BAL

Cytokines were measured using 497 Mesoscale Discovery. IL-16, IL-6, IL-10, IL-8, IL-2, IL-15, IL-12p40, CCL17 and MCP-1 were included in the V-PLEX Plus Proinflammatory Panel 1, V-PLEX Plus Chemokine Panel 1 and V-PLEX Plus Cytokine Panel 1 Human Kits (Meso Scale Diagnostics, LLC, Rockville, MD). The plate was read on a MESO Quickplex 500 SQ120 machine. Values below the limit of detection (LOD) were replaced with 50% of the LOD based on the lowest value of the standard curve for each analyte.

#### Lung Immunohistochemistry

Lung tissues sections were prepared as previously described^21^ from animals sacrificed 6 days post-challenge. Five micrometer sections of formalin-fixed, paraffin-embedded lung were incubated for 1 hour with the primary antibodies SARS-CoV-2 nucleoprotein (mouse IgG1, catalog number 40143-MM08, Sino Biological, Wayne, PA), Ionized calcium-binding adaptor (IBA1, rabbit polyclonal, catalog number 019-19741; Wako, Richmond, VA) or myeloperoxidase (MPO, rabbit polyclonal, catalog A0398, Agilent Dako, Santa Clara, CA) diluted in normal goat serum (NGS) at a concentration of 1:200, 1:100 and 1:6000 respectively. Secondary antibodies tagged with fluorochromes and diluted 1:1000 in NGS were incubated for 40 minutes. Slides were imaged with a digital slide scanner (Zeiss Axio Scan.Z1; Zeiss, White Plains, NY). Quantification of the stained cells were performed by laboratory staff who were blind of treatment allocation.

### Statistical analysis

Differences over the time on IgA, IgM, IgG and neutralizing antibody levels in the different vaccine formulations were assessed by repeated measures two-way ANOVA on log-transformed values. Kruskal-Wallis tests, followed by Dunn’s multiple comparisons post hoc analysis, were conducted independently to identify the significant differences of cell-mediated immune responses between vaccine formulations or timepoints. Kruskal-Wallis tests followed by Dunn’s multiple comparisons post hoc analysis were also performed to identify the significant differences between vaccine formulations in the cytokines and chemokines levels in BAL six days after challenge. The descriptive statistics and statistical comparisons were performed using GraphPad Prism software (Version 8.4.2; GraphPad Software, La Jolla, CA). Differences were declared statistically significant if *p*<0.05.

## Results

### Safety (pre-challenge)

None of the vaccine formulations induced any increase of body temperature during the seven days after the first or second immunization. All animals, including those in the Control group, lost weight during that study period after the first immunization, independent of the treatment. No weight loss was observed after the second immunization. After the first dose, one animal in the CoVLP group and another in the CoVLP+AS03 group developed mild (Grade 1) erythema at the injection site. Resorption of the erythema was observed within 2 to 3 days. No injection site observations were noted after the second dose.

After the first immunization, clinical observations were limited to reduced appetite (also seen in the Control group) and loose or soft stool, that occurred concurrently with normal stool. After the second immunization, no clinical signs were observed.

Hematology results did not reveal any safety concern up to 7 days after the first and second immunizations. The CoVLP+AS03 led to a transient mobilization of immune cells in the periphery one day after the first and second doses. This was mainly manifested by an increase of neutrophils and monocytes after the first immunization, and an increase of neutrophils after the second immunization, and was accompanied by a transient CRP signal reflecting the inflammatory reaction upon vaccination with the AS03-adjuvanted CoVLP. Immune cell mobilization and CRP increase were resolved 3 days after immunization and was not observed with unadjuvanted CoVLP or CoVLP+CpG.

### Immunogenicity

#### IgG, IgA and IgM

The RBD specific immunoglobulin (Ig) G, IgA and IgM were measured in pre-immune samples and at D14, D21 or D28 after the first immunization as well as 14 and 21 days after the second immunization (D42 and D49 respectively). No statistically significant differences were observed between the RBD specific IgG, IgA or IgM responses at D21 and at D28. Therefore, those two timepoints were combined (Figure 2). Unadjuvanted CoVLP induced a significant increase of RBD-specific IgG as early as 14 days after the first immunization (Figure 2A) that was maintained at 21-28 days. Both CpG and AS03 significantly increased RBD-specific IgG 14 and 21-28 days after immunization compared to unadjuvanted CoVLP, with a significantly greater impact of AS03 compared to CpG. All groups benefited from the boost sustaining the higher responses of adjuvanted formulations as compared to unadjuvanted CoVLP after the second dose (Figure 2A). The superior response for CoVLP+AS03 was maintained at all timepoints after the boost. Animals immunized with 2 doses of CoVLP+CpG experienced a small, although significant decrease in RBD-specific IgG levels between D42 and D49 (Figure 2A). Serum RBD-specific IgA responses were similar to RDB-specific IgG responses in intensity and ranking between groups (Figure 2B). Titres rose slightly in response to the unadjuvanted CoVLP but only barely reached statistical significance above baseline values after either the first or second dose. Animals that received adjuvanted CoVLP mounted stronger IgA responses that rose significantly at D14 and further increased with the second dose. Again, the IgA responses were consistently highest in the CoVLP+AS03 group at all timepoints. As observed with IgG, animals immunized with 2 doses of CoVLP+CpG experienced a small but significant decrease in RBD-specific IgA levels between D42 and D49 (Figure 2B).

**Figure 2:**
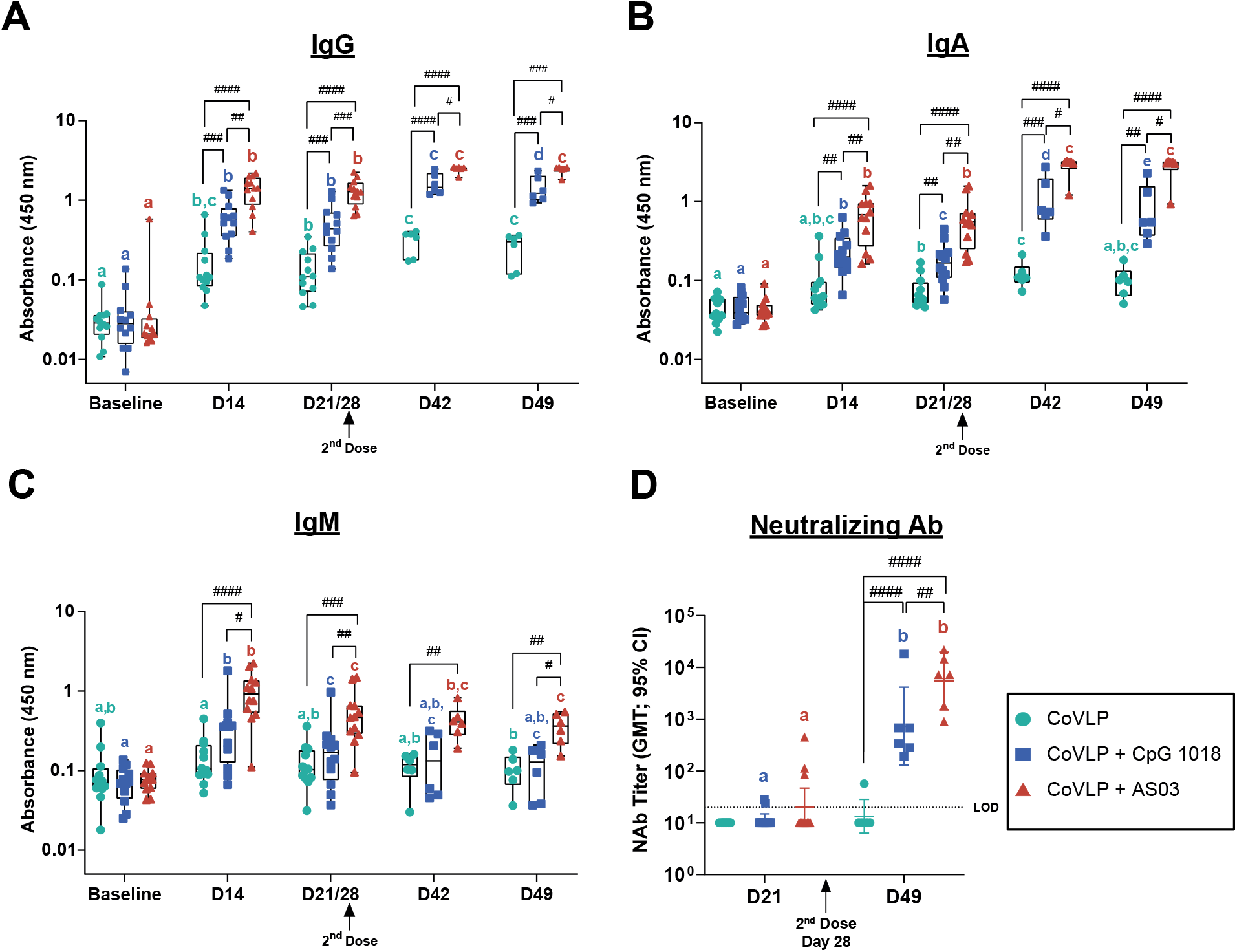
Humoral immune response. Anti receptor binding domain (RBD) IgG (**A**), IgA (**B**), IgM (**C**) and neutralizing antibody (**D**) responses in serum of macaques after the first and the second immunization, 28 days apart, with 15 μg CoVLP with and without CpG 1018 or AS03. Geometric mean (GMT) and 95% CI are represented. Different color-matched letters indicate significant differences between timepoints for each group (*p*<0.05). Significant differences between groups at each timepoints are indicated by # (^#^*p*<0.05, ^##^*p*<0.01, ^###^*p*<0.001, ^####^*p*<0.0001). Results from two-way ANOVA on log-transformed OD (450 nm) values (GraphPad Prism v8, San Diego, USA).

Although RBD-IgM levels remained significantly above baseline at all timepoints in the CoVLP+AS03 group, levels tended to fall in all groups at later times after the first dose and were not greatly influenced by the booster dose (Figure 2C), consistent with immunoglobulin class switching and the normal kinetics of IgM production following vaccination. RBD-specific IgM levels also rose in response to vaccination in CoVLP+CpG group but only reached statistical significance transiently at D14 and D21 (Figure 2C).

#### Neutralizing antibody

Serum NAb titers were measured at D21 and D49. None of the animals in the CoVLP group mounted a detectable NAb response after the first dose and NAb were detected in only 1/6 animals after the second dose. Among the animals that received adjuvanted vaccine only 2/12 and 3/12 in the CoVLP+CpG and CoVLP+AS03 groups respectively, mounted detectable responses after the first dose (Figure 2D). Responses after the second dose were significantly above baseline values in both the CoVLP+CpG and CoVLP+AS03 groups illustrating the significant impact on both adjuvants on the NAb response although levels in the CoVLP+AS03 group after the second dose were significantly higher than in the CoVLP+CpG group.

#### Cell-mediated immune response

The cell-mediated immune (CMI) response was measured in PBMC re-stimulated *ex vivo* with an S protein peptide pool at baseline, 7 and 21 days after the first and the second IM immunizations (D7, D21, D35 and D49 respectively). A single dose of 15 μg CoVLP+AS03 induced a significant increase of Ag-specific IL-2+ CD4 T cells in PBMC as soon as 7 days after the immunization with a clear boost effect (Figure 3A). Ag-specific IL-2+ CD4 T cells also significantly increased from baseline after the second dose of CoVLP+CpG. The higher response elicited by CoVLP+AS03 resulted in statistically significant differences between that group and both the unadjuvanted CoVLP or CoVLP+CpG groups at all timepoints with the exception of CoVLP+CpG at Day 49. No significant differences in Ag-specific IL-2+ CD4 T cells responses were observed between unadjuvanted CoVLP and CoVLP+CpG (Figure 3A). Adjuvanted CoVLP also increased polyfunctional CD4 T cells including triple positive Th1 (IL-2+ IFN-γ+ TNF-α+) CD4 T cells (Figure 3B). While this increase only reached statistical significance 21 days after the boost (D49) in the CoVLP+CpG group, a significant increase from baseline was detected as soon as 7 days after the first dose in animals vaccinated with CoVLP+AS03. Significantly higher levels of triple positive Th1 CD4 T cells as compared to baseline were maintained in the AS03 adjuvanted group at all timepoints (Figure 3B). However, due to the impact of the second dose of CoVLP+CpG on the quality of the CD4 T cell response (detailed hereafter), similar levels of triple positive Th1 CD4 T cells were observed in both adjuvanted formulations after the boost. An increase of CD4 T cells expressing CD40L was also observed, particularly in the CoVLP+AS03 group (Figure 3C) and most of the polyfunctional Th1 CD4 T, including the triple positive, expressed CD40L.

**Figure 3:**
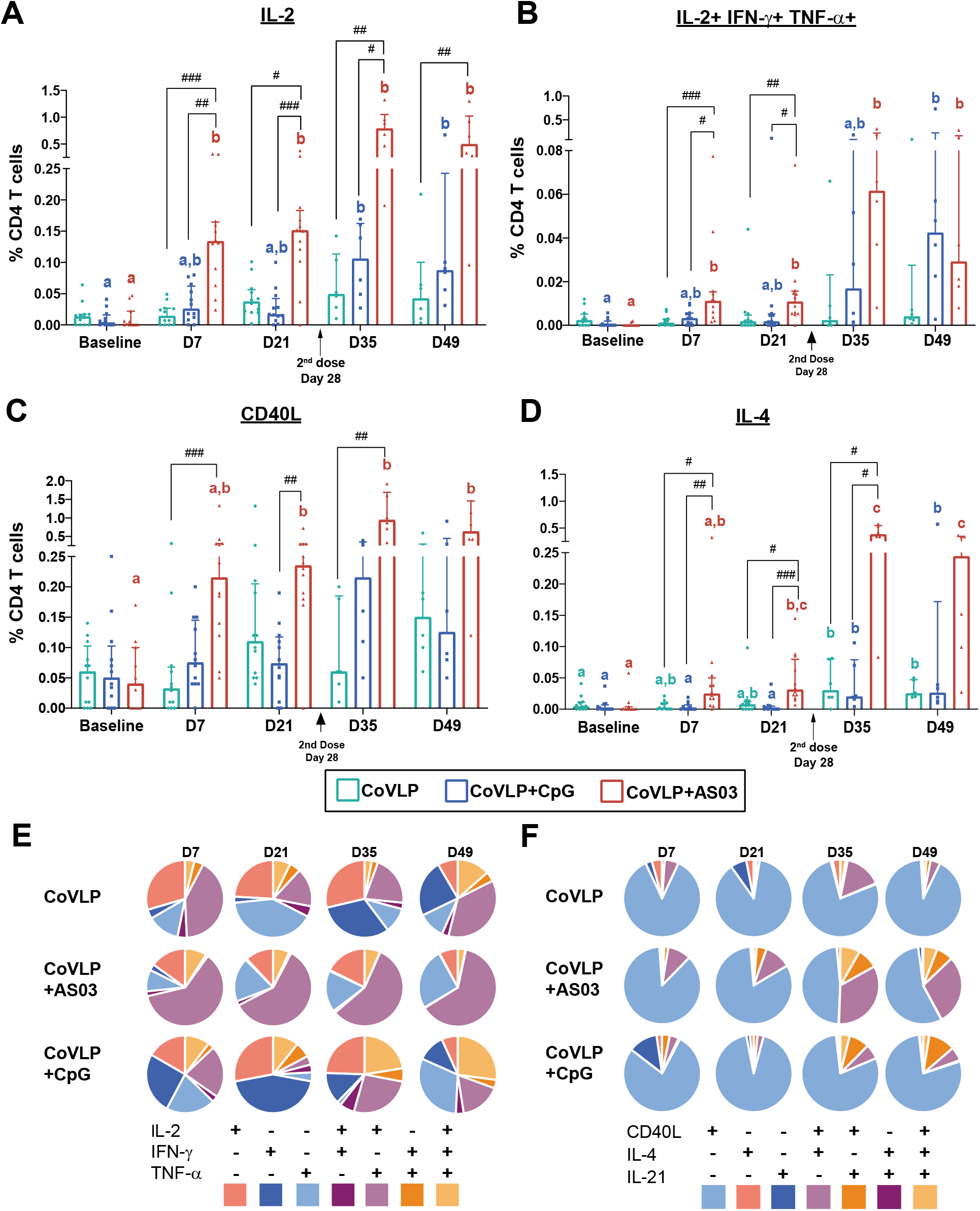
Cell-mediated immune response in PBMC of macaques at baseline, 7 and 21 days after the first and the second IM immunization with 15 μg CoVLP with and without CpG 1018 or AS03. Percent of S protein specific IL-2+ (**A**), triple posititive IL-2+ IFN-γ+ TNF-α+ (B), CD40L+ (**C**) and IL-4 (**D**) CD4 T cells. Median with interquartile range are represented. Within each treatment group, significant differences between timepoints were annotated as letters, a same letter on two bars indicates that no significant difference was detected between two timepoints. Differences were declared statistically significant if *p*<0.05. Significant differences between groups at each timepoint are indicated by # (^#^*p*<0.05, ^##^*p*<0.01, ^###^*p*<0.001) Kruskal-Wallis test with the Dunn post-hoc multiple comparisons test (GraphPad Prism v8.4.2, San Diego, USA). Qualitative distribution of Th1 (**E**) and Th2 (**F**) CD4 T cells according to the expressing of IL-2/IFN-γ/TNF-α or CD40L/IL-4/IL-21 respectively.

Significant induction of Ag-specific IL-4+ was observed after the boost in all groups, particularly in NHP immunized with CoVLP+AS03 (Figure 3D). However, the frequencies of Ag-specific IL-4+ CD4 T cells remained lower than Ag-specific IL-2+ CD4 T cells, even in the AS03 adjuvanted group.

Qualitative differences were observed in the CD4 T cells response elicited by the different vaccine regimens (Figure 3E&F). The response induced by CoVLP+AS03 was characterized by the higher frequency of IL-2+ TNF-α+ (IFN-γ-) CD4 T cells and remained relatively consistent with minimal impact of the boost on the relative proportion of the subpopulations based on the expressing of Th1 cytokines (Figure 3E).

In contrast, the quality of the Th1 response was strongly impacted by the boost in unadjuvanted CoVLP and CoVLP+CpG characterized by a predominance of monovalent CD4 T cells before boost and emergence of polyfunctionality after the boost. Notably, the relative proportion of triple positive IL-2+ IFN-γ+ TNF-α+ CD4 T cells increased after the second dose of CoVLP+CpG (Figure 3E).

The Ag-specific IL-4+ CD4 T cell response was strongly impacted by the boost with CoVLP+AS03 with emergence of IL-4+ IL-21+ CD4 T cells after the second dose (Figure 3F). The Th2 response in the CoVLP and CoVLP+CpG groups were characterized by the dominance of monovalent CD40L+ CD4 T cells (Figure 3F).

### Protection

In agreement with previously reported SARS-CoV-2 virus challenge in rhesus macaques, infection only resulted in mild symptoms in the control animals ^22,23^ with no significant differences in clinical scores between groups (Suppl. Figure 1). Animals in all groups experienced minor and transient reductions of peripheral oxygen saturation 6 days postchallenge; and slightly increased effort of breathing was noted in all groups, except the animals that received CoVLP+CpG. Soft stool, accompanied by liquid stool on some occasions, was observed after challenge in all groups and resolved by 20 days postchallenge, except in the unadjuvanted CoVLP group. Slight, although not statistically significant, increases of temperature in the Control animals were observed on day 6 and day 13 post-challenge (Suppl. Tables 3&4). A slight decrease in red blood cell distribution width (RDW) and an increase in mean platelet volume (MPV) were noted in all treatment groups following infection. In animals challenged after two doses of CoVLP, slight increases in leukocytes and neutrophils counts were noted in all treatment groups. On day 6 post-challenge, a transient increase in monocytes was observed in all experimental groups compared to pre-challenge levels. On day 13 post-challenge, monocyte counts were similar to those before challenge. On days 6 and 13 post-challenge, an increase in the number of lymphocytes was noted for animals treated with CoVLP alone or formulated with AS03. On day 20 post-challenge, lymphocyte levels of all animals were similar to pre-challenge values (Suppl. Tables 5&6).

#### Thoracic radiographs

Thoracic radiographs were performed 6, 13 and 20 days post-challenge. Not surprisingly given the mild clinical course in the challenged animals, radiographic changes and observations associated with infection were very subtle and limited.

<u>In animals challenged after one dose</u>, no abnormal observations were reported on say 6 after infection in animals immunized with unadjuvanted CoVLP. Tracheobronchial lymph node enlargements were reported on one lung of 2/6 animals in CoVLP+CpG group and loss of definition in the right caudal lung field was reported in 1/6 animal in the CoVLP+AS03 group (Table 1). A subtle increase in pulmonary opacity was observed at all time points after infection in one animal vaccinated with CoVLP+AS03 (Table 1). The incidence of radiographic findings was slightly higher in the Control group (1/4 animals 6 days after infection, 2/2 animals 13 days after infection and 2/2 animals 20 days after infection).

**Table 1:**
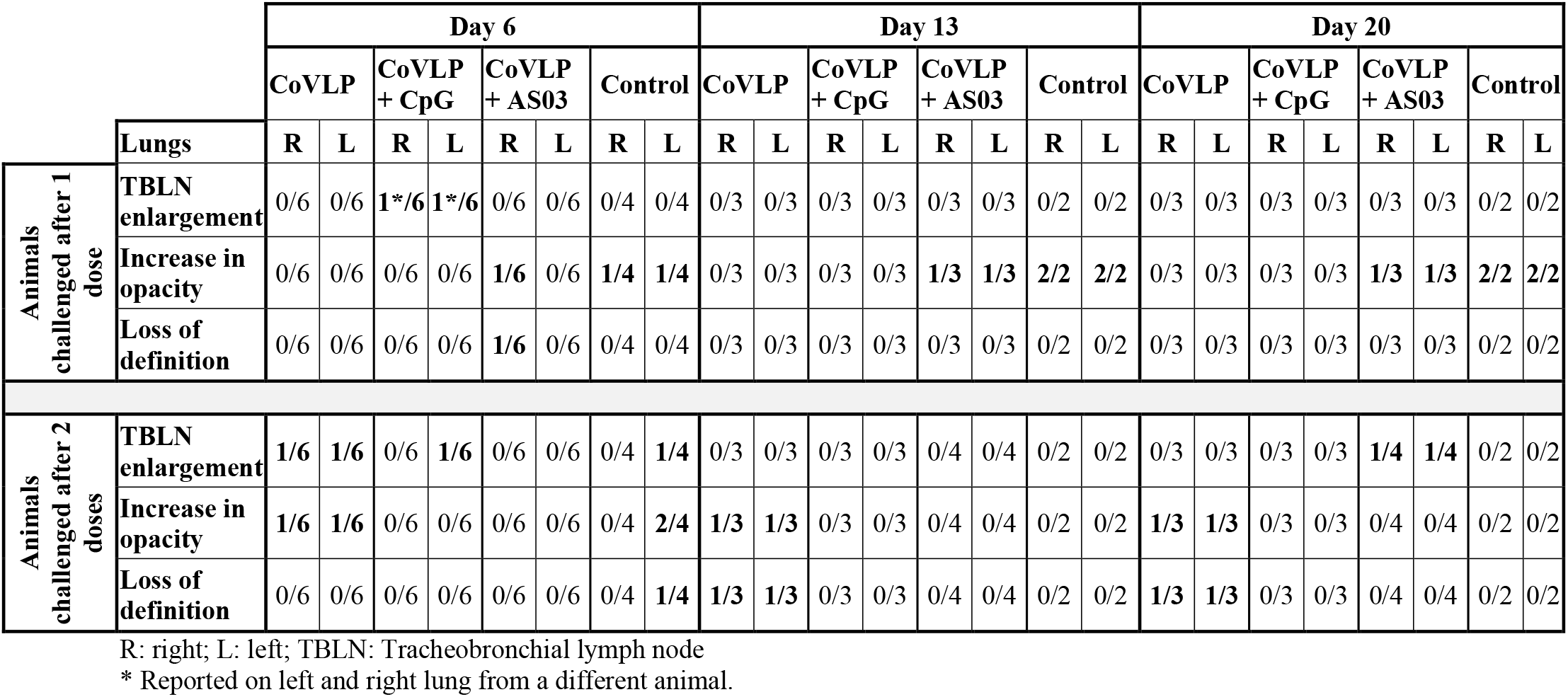
Thoracic Radiograph Observations and Incidence in Rhesus Macaques Infected with SARS-CoV-2 after One or Two Immunizations with Unadjuvanted CoVLP, CoVLP+CpG or CoVLP+AS03

<u>In animals challenged after two doses</u>, tracheobronchial lymph node enlargement was observed in CoVLP, CoVLP+CpG and Control groups (1/6, 1/6 and 1/4 animals, respectively) at 6 days post-challenge, and in 1/4 animal from the CoVLP+AS03 group, 20 days after challenge. Increase of pulmonary opacity was observed in one animal (in both lungs) from the CoVLP group at each timepoint (1/6, 1/3 and 1/3, respectively) and in 2/4 animals from the Control group, 6 days after the infection. Loss of lung definition was observed in 1/4 animal in the Control group 6 days post-challenge and in 1/3 animal in CoVLP group, 13 and 20 days after infection. At least one finding in one lung was observed in 37% (6 days post-challenge), 50% (13 days post-challenge) and 50% (20 days postchallenge) in the unvaccinated Control animals (Table 1). Radiographic findings were lower overall in the vaccinated groups: 17% (6 days post-challenge), 33% (13 days postchallenge) and 33% (20 days post-challenge) in the CoVLP group and even lower in animals that received two doses of an adjuvanted formulation i.e. 17% (6 days postchallenge), 0% (13 days post-challenge), 0% (20 days post-challenge) in the CoVLP+CpG group and 0% (6 days post-challenge), 0% (13 days post-challenge), 25% (20 days postchallenge) in the CoVLP+AS03 group.

Although these observations are limited by the small number of animals in each group and the modest pathogenicity of SARS-CoV-2 in rhesus macaques, the lower occurrence of lung findings in the vaccinated NHP after challenge suggests that two doses of the adjuvanted CoVLP may have provided a degree of protection in the lower respiratory tract. These observations are also notable for the absence of any suggestion of vaccine-enhanced disease.

#### Viral replication in swabs and BAL

Viral loads in the nasal swabs (sgRNA for the E gene) from the 8 Control animals at 6 days after challenge were highly variable (range undetectable to 10^8^ EqVC/mL). Although this variability and the relatively small number of animals in each treatment group limited our ability to perform statistical analyses, vaccination with one dose of unadjuvanted CoVLP had no obvious effect on virus load in the nasal secretions but a single dose of either adjuvanted CoVLP formulation appeared to reduce the median viral load by approximately 2 orders of magnitude with undetectable sgRNA levels in nasal swabs in 8/12 (66.6%) animals compared to 2/4 (50%) in the Control group (Figure 4A). A two-log reduction in nasal viral load was also observed in animals vaccinated with two doses of either CoVLP alone or CoVLP+CpG compared to the controls (Figure 4A). An even greater 4-log reduction in the median sgRNA level was seen in the NHP vaccinated with 2 doses of CoVLP+AS03. In this group, sgRNA was only detected in 1/6 animals (16.6%) compared to 3/4 (75%) of the Control group. No sgRNA was detected in pharyngeal swabs after 2 immunizations except in one Control animal (data not shown). SgRNA was detected from rectal swabs of three animals vaccinated with 2 doses of CoVLP+CpG and from the only animal of the CoVLP+AS03 group that also had detectable viral replication in nasal swab (data not shown). No sgRNA were detected in swab samples 13 and 20 days post-infection, with the exemption of one animal that received two doses of CoVLP alone (data not shown). SgRNA (E gene) in BAL was only detected in 3 vaccinated macaques 6 days after challenge (one immunized with a single dose of CoVLP+CpG, one with two doses of CoVLP+CpG and one with 2 doses CoVLP) and 3 control animals (data not shown). A more sensitive PCR assay measuring the *N* gene product^24^ was therefore used to measure viral replication in BAL (Figure 4B). In animals immunized with one dose, sg *N* RNA was not detected in the CoVLP alone group or in the animals that received CoVLP+AS03. Viral replication was observed in BAL from 3/6 (50%) and 2/4 (50%) animals in the CoVLP+CpG and Control groups, respectively (Figure 4B). In animals that received two doses, SgRNA was only detected in only 1/6 (16.6%) animal in the CoVLP+AS03 group compared to 5/6 (83.3%) of the animals in the CoVLP alone group, 2/6 (33.3%) in the CoVLP+CpG group and 2/4 Control animals (Figure 4B). Neither *E* nor *N* gene sgRNA was detected 13 or 20 days after challenge in BAL.

**Figure 4:**
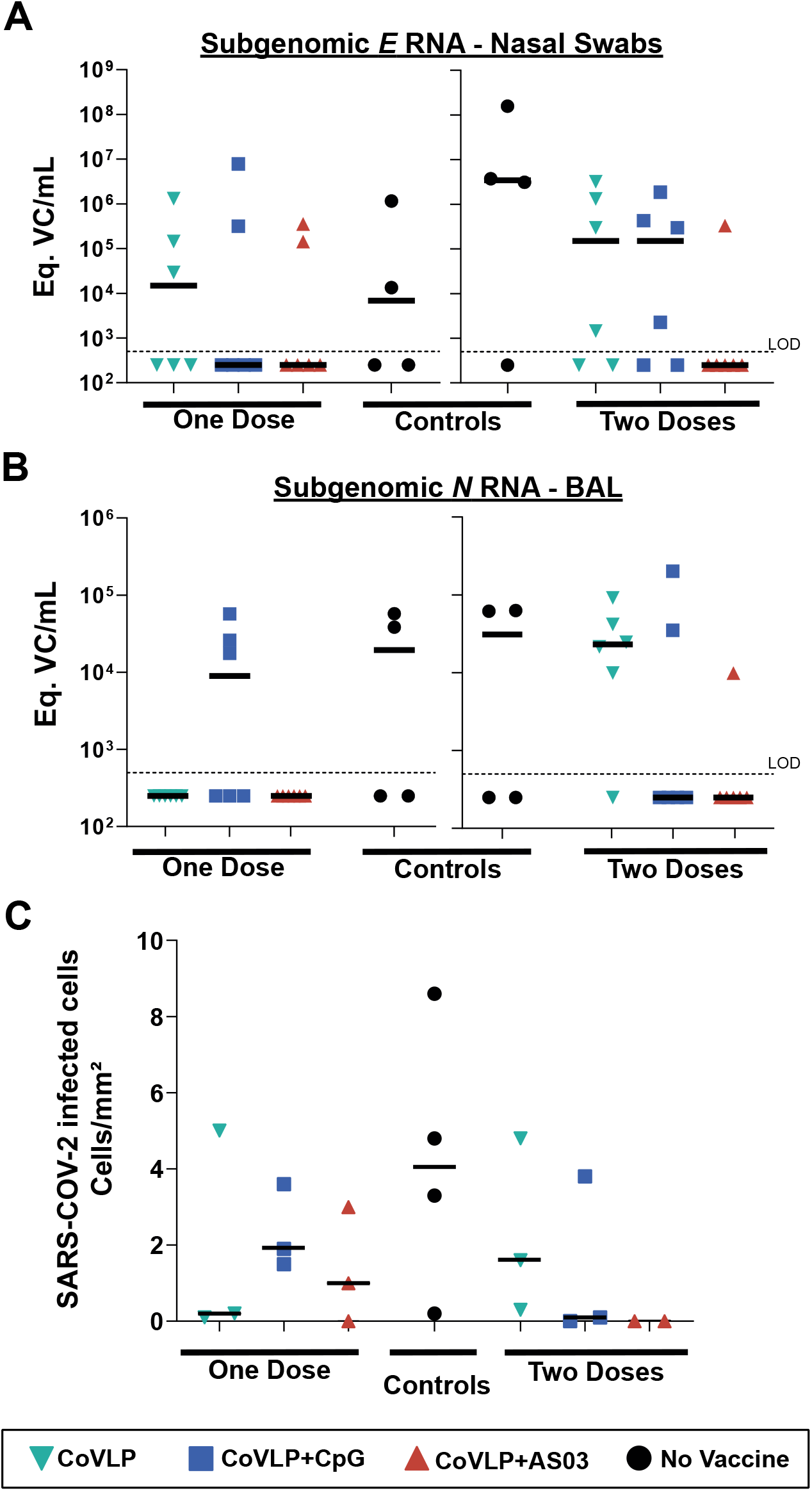
SARS-CoV-2 infection in lung 6 days post-challenge in rhesus macaques vaccinated with one or two dose(s) of CoVLP, CoVLP+CpG or CoVLP+AS03. Equivalent (Eq.) viral copy (VC) of subgenomic *E* gene RNA in nasal swabs (**A**) and subgenomic *N* gene RNA BAL (**B**) were measured by real time PCR. The dotted line represents the limit of detection of the method. Samples below the limit of detection were assigned a value of 250 Eq. VC/mL for graphical representation. SARS-CoV-2 infected cells measured by immunohistochemistry (**C**). Individual data and median (line) are represented.

#### SARS-CoV-2 infected cells and immune cell infiltration in lungs

SARS-CoV-2 infected cells and immune cell infiltration were detected by immunohistochemistry (Suppl. Figure 2A&B) and quantified in lung sections 6 days post-challenge (Figure 4C). Compared to the Control group, fewer SARS-CoV-2 infected cells were observed in the lungs of most animals that had received at least 1 dose of CoVLP with or without adjuvant and no SARS-CoV-2 infected cells were detected in the animals immunized with 2 doses of CoVLP+AS03 (Figure 4C). The lungs of Control animals were also characterized by the presence of immune cell infiltrates and expansion of the alveolar interstitium and perivascular spaces (Suppl. Figure 2C&D). While vaccination with one dose of adjuvanted CoVLP had minor impact, immunization with 2 doses of CoVLP strongly reduced the presence of macrophages, neutrophils and T lymphocytes in lungs 6 days after infection (Figure 5). Animals receiving 2 doses of adjuvanted CoVLP had a 3-fold reduction in macrophages (Figure 5A), a 2-to 4-fold reduction in neutrophils (Figure 5B) and a 12-to 20-fold reduction T cell infiltration (Figure 5C) compared to Control animals.

**Figure 5:**
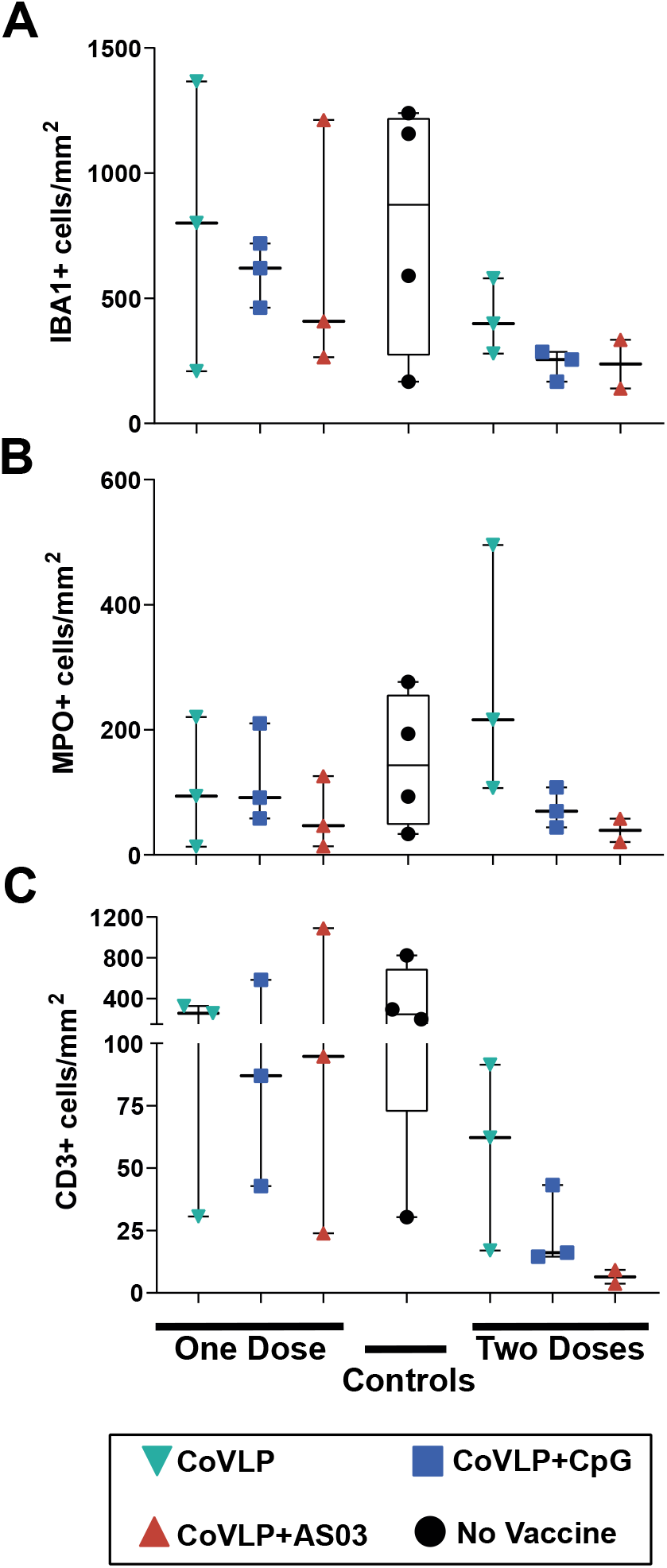
Immune cell infiltrations in lung sections from all animals 6 days after challenge. Macrophages (**A**, IBA1+ cells), neutrophils (B, MPO+ cells), and T lymphocytes (**C**, CD3+ cells) were quantified by immunohistochemistry. Individual data, median (line) and 75th/25th interquartile (box) are represented. Results are expressed as positive cells per mm^2^ of tissue.

#### Chemokines and cytokines in BAL

The most pronounced differences in cytokine and chemokine profiles in the BAL between treatment groups were observed 6 days post-challenge in the animals infected after 2 vaccine doses (Figure 6) while only limited differences were observed in animals challenged after one dose. In this context, it may be worth noting that the viral loads in nasal swabs from Control animals in the two-dose cohort were generally higher than in the Control group in the one-dose cohort. Although levels varied widely between animals in most groups, the Control animals tended to have the highest mean values for all of the cytokines/chemokines measured. Two doses of either unadjuvanted CoVLP or CoVLP+CpG had relatively little impact on the cytokine/chemokine profile except for lower IL-10 and IL-16 levels compared to the Control group (Figure 6). Although the difference only reached significance for IL-10, the animals that received two doses of CoVLP+AS03 generally had consistently but marginally lower levels of most pro-inflammatory cytokines and chemotactic factors in the BAL compared to the Control group (Figure 6); a finding consistent with the radiologic and histopathologic evidence of reduced inflammation in the CoVLP+AS03 group.

**Figure 6:**
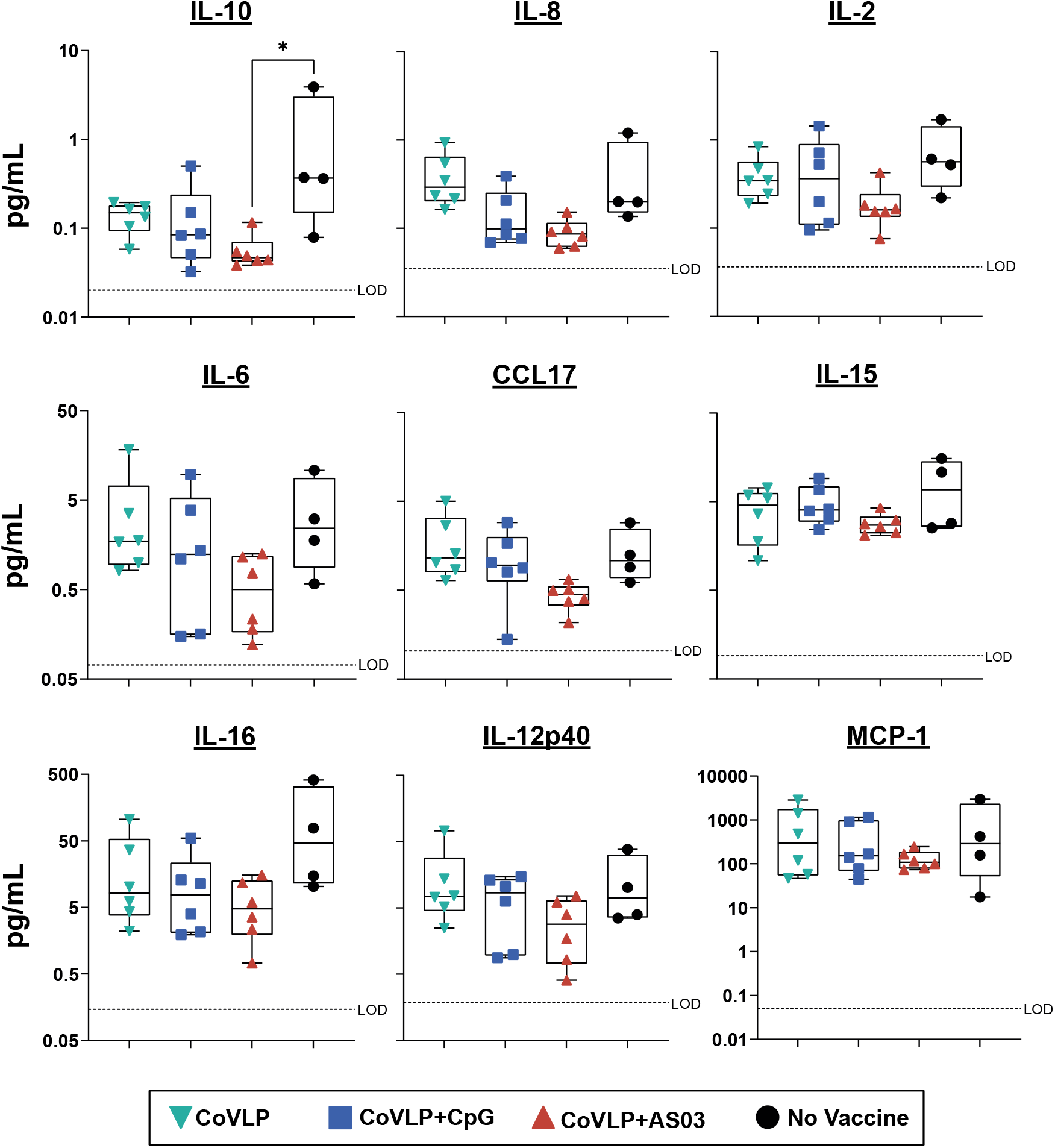
Cytokines and Chemokines in BAL samples. collected six days post-challenge with SARS-CoV-2 from rhesus macaques immunized with two doses of CoVLP, CoVLP+CpG or CoVLP+AS03. Dotted lines indicate the LOD for each cytokine/chemokine. Individual data, median (line) and 75th/25th interquartile (box) are represented. Statistical comparisons were performed using a non-parametric Kruskal-Wallis test. The Dunn post-hoc multiple comparisons test was used (\**p* < 0.05).

## Discussion

No safety issues were observed after one or two injections of either the adjuvanted or unadjuvanted CoVLP vaccine candidate. The transient evidence of inflammation and mobilization of neutrophils (and to a lesser extend monocytes) in the blood of the animals that received CoVLP+AS03 are consistent with the inflammatory response observed upon vaccination with AS03-adjuvanted vaccines^12,14^.

Although one dose of unadjuvanted CoVLP was sufficient to elicit anti-RBD IgG as soon as D14, both the addition of an adjuvant and the second dose significantly improved the humoral response. The beneficial effect of CpG 1018 and AS03 on the humoral response has been previously demonstrated for hepatitis B^25–27^ and influenza^16,17^ vaccines respectively. More recently, benefits of CpG 1018 (alone or with Alum) or AS03 on antibody response induced by COVID-19 vaccine candidates have been reported in NHP and humans^28–30^. As observed in the present study, the vaccine candidates adjuvanted with AS03 elicited higher humoral and T cell responses than the CpG 1018-adjuvanted formulations. Monocyte-derived cells, rather than bona-fide dendritic cells, seem to play an important role in adjuvanticity of AS03. In addition to recruitment and activation of cells at the site of injection, emulsions also favor the uptake of the antigen by antigenpresenting cells and its transport to the draining lymph node. The presence of α-tocopherol was associated with an increased uptake of antigen by monocytes possibly contributing to the higher antibody response^11^. The IgG and IgA induced after the first dose appeared to have a limited neutralizing potential although moderate protection was observed in animals vaccinated with one dose of adjuvanted CoVLP illustrating the potential role of nonneutralizing antibodies and cell-mediated immunity^31–33^. Levels of NAb induced after the second dose of CoVLP+AS03 have been reported to be protective in macaques following IgG adoptive transfer^34^.

CoVLP+AS03 also elicited a strong IL-2 driven CD4 response with a significant increase of polyfunctional including triple positive Ag-specific IL-2+ IFN-γ+ TNF-α+ Th1 positive CD4 T cells as soon as 7 days after the prime. Similar to the response induced by AS03 adjuvanted inactivated split H5N1 influenza vaccine in humans^12,17^, the S protein-specific CD4 T cells induced by CoVLP+AS03 were also characterized by the expression of CD40L. As expected, the CpG adjuvanted formulation induced a more Th1-biased response characterized by a higher relative proportion of Ag-specific IFN-γ+ CD4 T cells as compared to CoVLP+AS03. The response elicited by CoVLP+CpG was characterized by monovalent IL-2 and IFN-γ CD4 T cells after the first dose and the emergence of polyfunctional CD4 T cells after the second dose with minor induction of S protein-specific IL-4+ CD4 T cells.

Human COVID-19 infection results in a Th1 response and a large proportion of the SARS-CoV-2-specific CD4+ T cells in the peripheral blood samples of recovered patients were directed against S protein epitopes^35,36^. Studies of acute and convalescent COVID-19 patients have observed that T cell responses are associated with reduced disease^37,38^, suggesting that SARS-CoV-2-specific CD4+ and CD8+ T cell responses contribute significantly to control and resolution of the infection. Studies in mouse models generally confirm the critical role of T cells in viral clearance during coronavirus infections^39–41^. Moreover, polyfunctional Th1 cells that target viral antigens have been demonstrated to provide protection against other viral infections^42–44^ supporting the hypothesis that S-specific CD4 T cells contribute to the observed reduction of viral load in animals immunized with the adjuvanted CoVLP vaccine candidate. Additionally, it has been recently reported that sequences of the vast majority of SARS-CoV-2 T cell epitopes, including epitopes in the S protein, are not affected by the mutations found in the predominant variants ^45^ in agreement with potential important role of cell-mediated immune response in cross-protection against the emerging variants. The Th1 response elicited by CoVLP+AS03 was characterized by the predominance of IL-2+, TNF-α+ (IFN-γ-) CD4 T cells. Influenza-induced IL-2+ TNF-α+ (IFN-γ-) CD4 T cells have been described as uncommitted Th1 primed precursor (Thpp) preferentially promoted by newly encountered antigen whereas common, multiply-boosted influenza epitope-specific CD4 T cells expressed IFN-γ after infections^46,47^. We previously reported the induction of Thpp after prime-boost immunizations (21 days apart) with VLP presenting a H5N1 influenza hemagglutinin protein adjuvanted with either Alum or glucopyranosyl lipid adjuvant-stable emulsion (GLA-SE). Induction of Thpp CD4 T cells was also observed, although on a lower relative proportion, after prime-boost immunization 21 days apart with subunit vaccine consisting in the SARS-CoV-2 RBD domain displayed on a two-component protein nanoparticle adjuvanted with AS03 or Alum^29^. Thpp have been proposed to serve as a reservoir of memory CD4+ T cells with effector potential^47^. As mentioned above, CoVLP+AS03 immunization was also associated with a higher humoral response. By secreting IL-4, Th2 cells can instruct B cells to produce IgG1 and IgE. The higher induction of CD40L IL-4+/IL-21+ observed in the CoVLP+AS03 group is consistent with the higher antibody response measured in those animals. Co-expression of CD40L+ and IL-21+ can be indicative of the T follicular helper cell (Tfh) engagement which play a critical role in B-cell maturation and the establishment of memory^48,49^.

Although challenged with a relatively high viral dose, the absence of major clinical signs is in agreement with previously reported challenges in that animal model^23,29,34,50^.

Levels of sgRNA in placebo control animals are higher or in agreement with previously reported values in NHP after SARS-CoV-2 challenge^34,50^. The reduced viral replication in nasal swabs, the lower incidence of clinical observations on thoracic radiographs and the reduced SARS-CoV-2 infected cell in lungs especially in animals vaccinated with two doses of CoVLP+AS03 indicated the potential protection induced by the adjuvanted CoVLP along with a reduction of viral shedding.

High levels of pro-inflammatory cytokine IL-6 and IL-10, have been associated with severe outcomes during COVID-19 infection (e.g.: hospitalization, Intensive Care Unit admission)^51,52^, and low levels of CCL17 have been associated with lower viral replication in rhesus macaques, while high levels correlate with higher viral load in bronchoalveolar brush samples ^21^. Not surprisingly, the lower level of pro-inflammatory cytokines and chemotactic factors was associated with less infiltration of immune cells in the lungs of animals immunized with two doses of the adjuvanted CoVLP. Although migration of immune cells to the site of infection is essential to resolve viral infection, persistence of lymphoid cell infiltrates have been associated with uncontrolled inflammation and lung damage in coronavirus infections, including COVID-19^53–56^. The nominal presence of immune cells associated with lower levels of pro-inflammatory cytokines and chemotactic factors in BAL of vaccinated animals contributed to demonstrate the beneficial effect of two doses of adjuvanted vaccine while concurring to demonstrate the absence of VAED.

The observed benefits provided by AS03 on CoVLP were consistent with reported increase immunogenicity and protection^28,29,57^. The detailed CMI response reported in the present NHP study, including the possible engagement of Tfh suggest a possible beneficial effect of AS03 on the durability of the immune response. Based on this study and the results from the Phase 1 clinical trial^30^, CoVLP+AS03 was selected for the Phase 2/3 studies.

## Acknowledgements

The authors would like to acknowledge Robbert Van Der Most, Cindy Gutzeit, Philipe Boutet, Ulrike Krauze, Ellen Oe, Walthère Dewé, Margherita Coccia and Eric Destexhe from GSK as well as Dong Yu, Darren Campbell and Paula Traquina from Dynavax for critical review of the manuscript. The authors also wish to acknowledge all the Medicago employees and their contractors for their exceptional dedication and professionalism.

## References

1. Ou, X., et al. Characterization of spike glycoprotein of SARS-CoV-2 on virus entry and its immune cross-reactivity with SARS-CoV. Nat Commun 11, 1620 (2020).

2. Li, F. Structure, Function, and Evolution of Coronavirus Spike Proteins. Annu Rev Virol 3, 237–261 (2016).

3. Grant, O.C., Montgomery, D., Ito, K. & Woods, R.J. Analysis of the SARS-CoV-2 spike protein glycan shield reveals implications for immune recognition. Sci Rep 10, 14991 (2020).

4. Hoffmann, M., Kleine-Weber, H. & Pohlmann, S. A Multibasic Cleavage Site in the Spike Protein of SARS-CoV-2 Is Essential for Infection of Human Lung Cells. Mol Cell 78, 779–784 e775 (2020).

5. D’Aoust, M.A., et al. The production of hemagglutinin-based virus-like particles in plants: a rapid, efficient and safe response to pandemic influenza. Plant biotechnology journal 8, 607–619 (2010).

6. Du, L., et al. The spike protein of SARS-CoV--a target for vaccine and therapeutic development. Nat Rev Microbiol 7, 226–236 (2009).

7. Pallesen, J., et al. Immunogenicity and structures of a rationally designed prefusion MERS-CoV spike antigen. Proc Natl Acad Sci U S A 114, E7348–E7357 (2017).

8. Del Giudice, G., Rappuoli, R. & Didierlaurent, A.M. Correlates of adjuvanticity: A review on adjuvants in licensed vaccines. Semin Immunol 39, 14–21 (2018).

9. Campbell, J.D. Development of the CpG Adjuvant 1018: A Case Study. Methods Mol Biol 1494, 15–27 (2017).

10. Shirota, H. & Klinman, D.M. Recent progress concerning CpG DNA and its use as a vaccine adjuvant. Expert Rev Vaccines 13, 299–312 (2014).

11. Morel, S., et al. Adjuvant System AS03 containing alpha-tocopherol modulates innate immune response and leads to improved adaptive immunity. Vaccine 29, 2461–2473 (2011).

12. Burny, W., et al. Different Adjuvants Induce Common Innate Pathways That Are Associated with Enhanced Adaptive Responses against a Model Antigen in Humans. Front Immunol 8, 943 (2017).

13. De Mot, L., et al. Transcriptional profiles of adjuvanted hepatitis B vaccines display variable interindividual homogeneity but a shared core signature. Sci Transl Med 12 (2020).

14. Howard, L.M., et al. AS03-Adjuvanted H5N1 Avian Influenza Vaccine Modulates Early Innate Immune Signatures in Human Peripheral Blood Mononuclear Cells. J Infect Dis 219, 1786–1798 (2019).

15. Khurana, S., et al. AS03-adjuvanted H5N1 vaccine promotes antibody diversity and affinity maturation, NAI titers, cross-clade H5N1 neutralization, but not H1N1 cross-subtype neutralization. NPJ Vaccines 3, 40 (2018).

16. Leroux-Roels, I., et al. Antigen sparing and cross-reactive immunity with an adjuvanted rH5N1 prototype pandemic influenza vaccine: a randomised controlled trial. Lancet 370, 580–589 (2007).

17. Moris, P., et al. H5N1 influenza vaccine formulated with AS03 A induces strong cross-reactive and polyfunctional CD4 T-cell responses. Journal of clinical immunology 31, 443–454 (2011).

18. Cohet, C., et al. Safety of AS03-adjuvanted influenza vaccines: A review of the evidence. Vaccine 37, 3006–3021 (2019).

19. Wrapp, D., et al. Cryo-EM structure of the 2019-nCoV spike in the prefusion conformation. Science 367, 1260–1263 (2020).

20. Roltgen, K., et al. Defining the features and duration of antibody responses to SARS-CoV-2 infection associated with disease severity and outcome. Sci Immunol 5 (2020).

21. Fahlberg, M.D., et al. Cellular events of acute, resolving or progressive COVID-19 in SARS-CoV-2 infected non-human primates. Nat Commun 11, 6078 (2020).

22. Blair, R.V., et al. Acute Respiratory Distress in Aged, SARS-CoV-2-Infected African Green Monkeys but Not Rhesus Macaques. Am J Pathol 191, 274–282 (2021).

23. Munster, V.J., et al. Respiratory disease in rhesus macaques inoculated with SARS-CoV-2. Nature 585, 268–272 (2020).

24. Kim, D., et al. The Architecture of SARS-CoV-2 Transcriptome. Cell 181, 914–921 e910 (2020).

25. Halperin, S.A., et al. Safety and immunogenicity of different two-dose regimens of an investigational hepatitis B vaccine (hepatitis B surface antigen co-administered with an immunostimulatory phosphorothioate oligodeoxyribonucleotide) in healthy young adults. Vaccine 30, 5445–5448 (2012).

26. Heyward, W.L., et al. Immunogenicity and safety of an investigational hepatitis B vaccine with a Toll-like receptor 9 agonist adjuvant (HBsAg-1018) compared to a licensed hepatitis B vaccine in healthy adults 40-70 years of age. Vaccine 31, 5300–5305 (2013).

27. Leroux-Roels, G., et al. Impact of adjuvants on CD4(+) T cell and B cell responses to a protein antigen vaccine: Results from a phase II, randomized, multicenter trial. Clin Immunol 169, 16–27 (2016).

28. Liang, J.G., et al. S-Trimer, a COVID-19 subunit vaccine candidate, induces protective immunity in nonhuman primates. Nat Commun 12, 1346 (2021).

29. Arunachalam, P.S., et al. Adjuvanting a subunit SARS-CoV-2 nanoparticle vaccine to induce protective immunity in non-human primates. bioRxiv, 2021.2002.2010.430696 (2021).

30. Ward, B.J., et al. Phase 1 Randomized Trial of a Plant-Derived Virus-Like Particle Vaccine for Covid-19. Nature Medicine (accepted for publication).

31. Tauzin, A., et al. A single BNT162b2 mRNA dose elicits antibodies with Fc-mediated effector functions and boost pre-existing humoral and T cell responses. bioRxiv (2021).

32. Yu, J., et al. DNA vaccine protection against SARS-CoV-2 in rhesus macaques. Science 369, 806–811 (2020).

33. Bartsch, Y.C., et al. Discrete SARS-CoV-2 antibody titers track with functional humoral stability. Nat Commun 12, 1018 (2021).

34. McMahan, K., et al. Correlates of protection against SARS-CoV-2 in rhesus macaques. Nature 590, 630–634 (2021).

35. Dan, J.M., et al. Immunological memory to SARS-CoV-2 assessed for up to 8 months after infection. Science 371 (2021).

36. Nelde, A., et al. SARS-CoV-2-derived peptides define heterologous and COVID-19-induced T cell recognition. Nat Immunol 22, 74–85 (2021).

37. Rydyznski Moderbacher, C., et al. Antigen-Specific Adaptive Immunity to SARS-CoV-2 in Acute COVID-19 and Associations with Age and Disease Severity. Cell 183, 996–1012 e1019 (2020).

38. Sette, A. & Crotty, S. Adaptive immunity to SARS-CoV-2 and COVID-19. Cell 184, 861–880 (2021).

39. Chen, J., et al. Cellular immune responses to severe acute respiratory syndrome coronavirus (SARS-CoV) infection in senescent BALB/c mice: CD4+ T cells are important in control of SARS-CoV infection. J Virol 84, 1289–1301 (2010).

40. Zhao, J., Zhao, J. & Perlman, S. T cell responses are required for protection from clinical disease and for virus clearance in severe acute respiratory syndrome coronavirus-infected mice. J Virol 84, 9318–9325 (2010).

41. Zhao, J., et al. Airway Memory CD4(+) T Cells Mediate Protective Immunity against Emerging Respiratory Coronaviruses. Immunity 44, 1379–1391 (2016).

42. Appay, V., van Lier, R.A., Sallusto, F. & Roederer, M. Phenotype and function of human T lymphocyte subsets: consensus and issues. Cytometry. Part A: the journal of the International Society for Analytical Cytology 73, 975–983 (2008).

43. Sant, A.J. & McMichael, A. Revealing the role of CD4(+) T cells in viral immunity. The Journal of experimental medicine 209, 1391–1395 (2012).

44. Seder, R.A., Darrah, P.A. & Roederer, M. T-cell quality in memory and protection: implications for vaccine design. Nature reviews. Immunology 8, 247–258 (2008).

45. Tarke, A., et al. Negligible impact of SARS-CoV-2 variants on CD4 (+) and CD8 (+) T cell reactivity in COVID-19 exposed donors and vaccinees. bioRxiv (2021).

46. Deng, N., Weaver, J.M. & Mosmann, T.R. Cytokine diversity in the Th1-dominated human anti-influenza response caused by variable cytokine expression by Th1 cells, and a minor population of uncommitted IL-2+IFNgamma-Thpp cells. PLoS One 9, e95986 (2014).

47. Weaver, J.M., et al. Increase in IFNgamma(-)IL-2(+) cells in recent human CD4 T cell responses to 2009 pandemic H1N1 influenza. PLoS One 8, e57275 (2013).

48. Crotty, S. Follicular helper CD4 T cells (TFH). Annu Rev Immunol 29, 621–663 (2011).

49. Zhu, J. T helper 2 (Th2) cell differentiation, type 2 innate lymphoid cell (ILC2) development and regulation of interleukin-4 (IL-4) and IL-13 production. Cytokine 75, 14–24 (2015).

50. Mercado, N.B., et al. Single-shot Ad26 vaccine protects against SARS-CoV-2 in rhesus macaques. Nature 586, 583–588 (2020).

51. McGonagle, D., Sharif, K., O’Regan, A. & Bridgewood, C. The Role of Cytokines including Interleukin-6 in COVID-19 induced Pneumonia and Macrophage Activation Syndrome-Like Disease. Autoimmun Rev 19, 102537 (2020).

52. Pandolfi, L., et al. Broncho-alveolar inflammation in COVID-19 patients: a correlation with clinical outcome. BMC Pulm Med 20, 301 (2020).

53. Ye, J., et al. Molecular pathology in the lungs of severe acute respiratory syndrome patients. Am J Pathol 170, 538–545 (2007).

54. Carsana, L., et al. Pulmonary post-mortem findings in a series of COVID-19 cases from northern Italy: a two-centre descriptive study. Lancet Infect Dis 20, 1135–1140 (2020).

55. Fox, S.E., et al. Pulmonary and cardiac pathology in African American patients with COVID-19: an autopsy series from New Orleans. Lancet Respir Med 8, 681-686 (2020).

56. Schurink, B., et al. Viral presence and immunopathology in patients with lethal COVID-19: a prospective autopsy cohort study. Lancet Microbe 1, e290–e299 (2020).

57. Richmond, P., et al. Safety and immunogenicity of S-Trimer (SCB-2019), a protein subunit vaccine candidate for COVID-19 in healthy adults: a phase 1, randomised, double-blind, placebo-controlled trial. Lancet 397, 682–694 (2021).

