## Supplementary information for "Safety, immunogenicity and protection provided by unadjuvanted and adjuvanted formulations of recombinant plant-derived virus-like particle vaccine candidate for COVID-19 in non-human primates"

### Supplemental Material

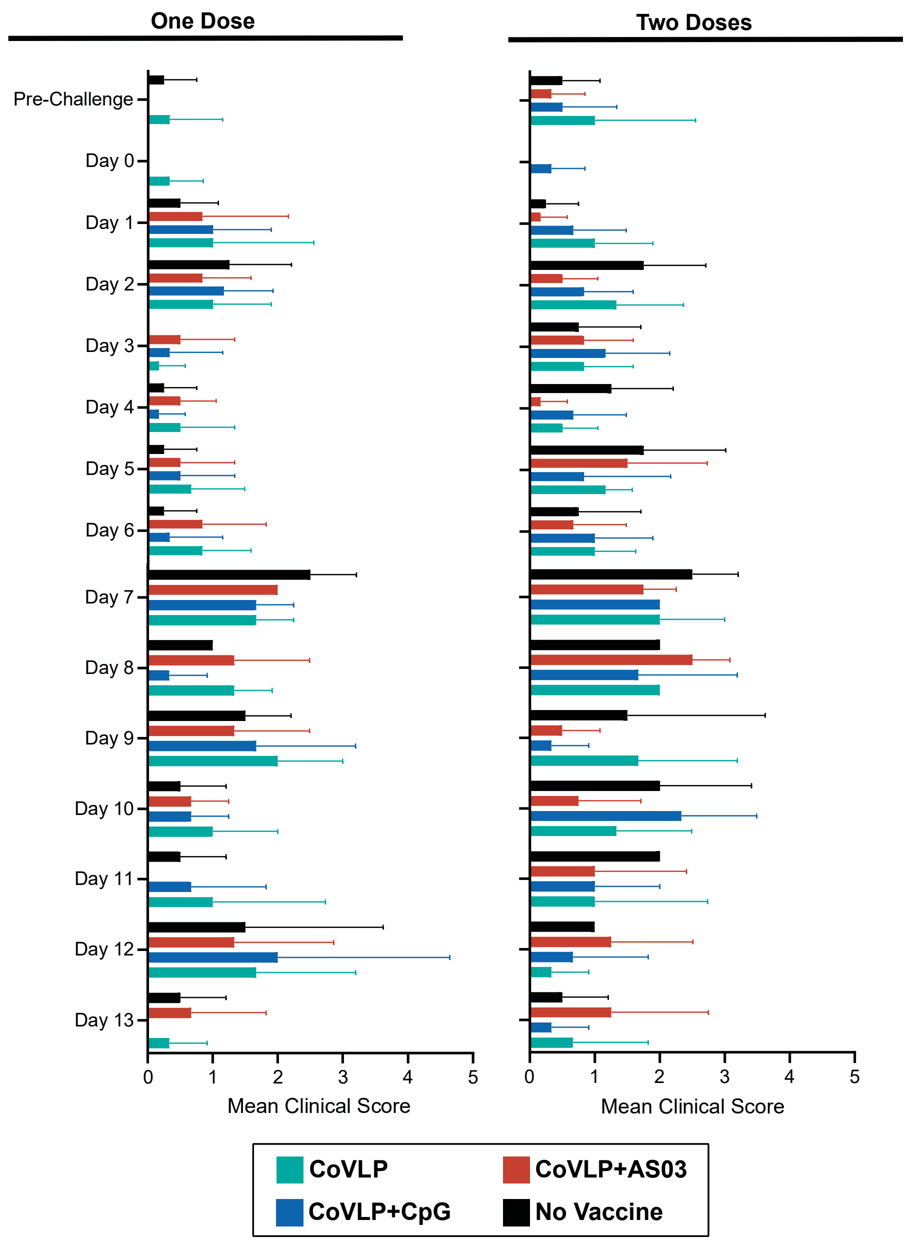

**Supplementary Figure 1: Global clinical score in rhesus macaques infected with SARS-CoV-2 after challenge** in animals immunized with one or two doses of CoVLP unadjuvanted or adjuvanted with AS03 or CpG 1018. Clinical observations were performed every day after challenge with SARS-CoV-2. Animals were individually scored for the severity of 6 different clinical observations (state of responsiveness, presence of discharge, skin condition, respiratory condition, food consumption and fecal condition). Global clinical score represents the addition of all clinical observations.

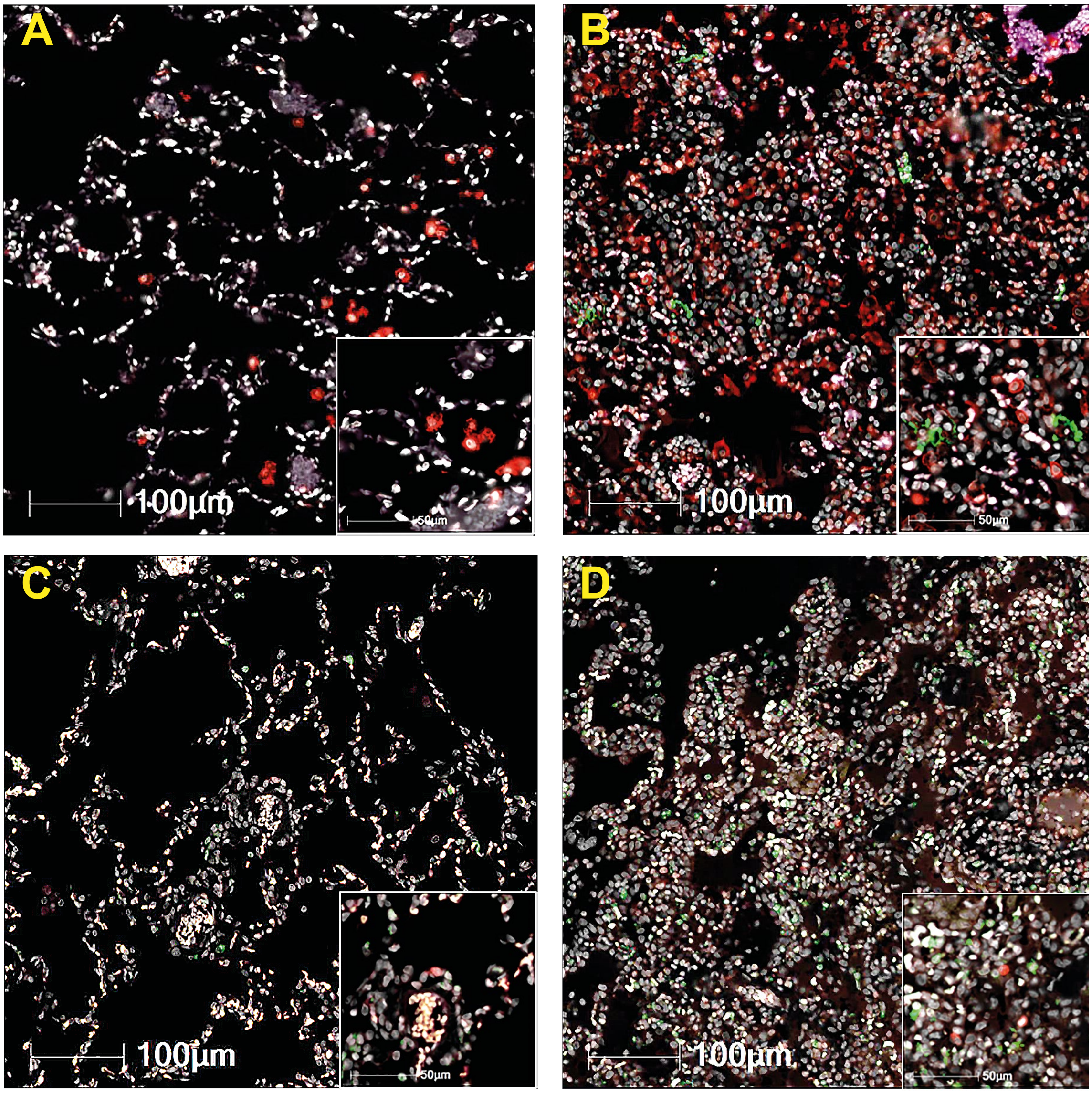

**Supplementary Figure 2: Fluorescent Immunohistochemistry of the Lung from SARS-CoV-2 Infected Rhesus Macaques 6 days Post-challenge**. Lung sections were stained for detection of macrophages (IBA1+, red) and SARS-CoV-2 nucleoprotein (green) (A and B) or T lymphocytes (CD3+, red) and neutrophils (MPO+, green) (C and D) in an animal with no detectable viral replication (A&C) or with high (>60,000 Eq. VC/mL) viral replication (B&D) in BAL.

**Supplementary Table 1: Treatment groups**

| Group No. | n | Group Description | CoVLP Dose (µg)[1] | Adjuvant Dose | Immunization (Days) | Challenge (Days) |
|---|---|---|---|---|---|---|
| 1 | 6 | CoVLP | 15 | 0 | 0 | 28 |
| 2 | 6 | CoVLP+CpG 1018 | 15 | 3 mg | 0 | 28 |
| 3 | 6 | CoVLP+AS03 | 15 | Ratio 1:1 (v:v) | 0 | 28 |
| 4 | 4 | Control (Cohort one dose) | -[(2)] | 0 | 0 | 28 |
| 5 | 6 | CoVLP | 15 | 0 | 0 and 28 | 57 |
| 6 | 6 | CoVLP + CpG 1018 | 15 | 3 mg | 0 and 28 | 57 |
| 7 | 6 | CoVLP + AS03 | 15 | Ratio 1:1 (v:v) | 0 and 28 | 57 |
| 8 | 4 | Control (Cohort two dose) | -[(2)] | 0 | 0 and 28 | 57 |

[1] Total CoVLP proteins corrected by purity.
[2] The no vaccine control animals were administered Phosphate Buffered Saline (PBS) Solution.

**Supplementary Table 2: Immunological markers used to assess the cell-mediated response by flow cytometry**

| | | Fluorochrome | Clone | Manufacturer |
|---|---|---|---|---|
| | Viability | Zombie UV fixable viability dye | | |
| Surface Stain | CD3 | BV650 | SP34-2 | BD Biosciences |
| | CD4 | BV605 | OKT4 | BioLegend |
| | CD8 | BUV563 | RPA-T8 | BD Biosciences |
| Intracellular Stain | CD154/CD40L | BV421 | 24-31 | BioLegend |
| | IL-2 | FITC | MQ1-17H12 | BioLegend |
| | IFN-γ | A700 | 4S.B3 | BioLegend |
| | TNF-α | PE-Cy7 | Mab11 | E-Bioscience |
| | IL-4 | PE | MP4-25D2 | BioLegend |
| | IL-21 | APC | 3A3-N2 | BioLegend |

#### Supplementary Table 3: Post-Challenge Clinical Signs in Rhesus Macaques Infected with SARS-CoV-2 after One Immunization with CoVLP Unadjuvanted or Adjuvanted with AS03 or CpG 1018 Adjuvants

| Parameter | | Pre-Challenge[1] | | | | Post-Challenge[2] | | | | | | | | | | | |
|---|---|---|---|---|---|---|---|---|---|---|---|---|---|---|---|---|---|
| | | | | | | Day 6 | | | | Day 13 | | | | Day 20 | | | |
| | | **CoVLP** | **CoVLP + CpG 1018** | **CoVLP + AS03** | **Control** | **CoVLP** | **CoVLP + CpG 1018** | **CoVLP + AS03** | **Control** | **CoVLP** | **CoVLP + CpG 1018** | **CoVLP + AS03** | **Control** | **CoVLP** | **CoVLP + CpG 1018** | **CoVLP + AS03** | **Control** |
| Temp. Change (°C)[3,4] | | 0.0±0.0 | 0.0±0.0 | 0.0±0.0 | 0.0±0.0 | 0.1±0.7 | 0.3±0.6 | 0.0±0.4 | 0.8±0.8 | 0.0±0.4 | 0.0±0.3 | 0.0±0.7 | 0.9±0.3 | -0.7±0.7 | -0.2±0.5 | -0.2±0.3 | 0.3±0.1 |
| $\delta$ Weight (%)[3,4] | | 100±0 | 100±0 | 100±0 | 100±0 | 102±3 | 103±3 | 103±2 | 102±2 | 101±1 | 103±2 | 103±1 | 106±0 | 104±3 | 107±2 | 103±3 | 109±1 |
| Respiratory Rate[3] | | 36±8 | 40±11 | 38±11 | 44±5 | 37±9 | 43±10 | 38±6 | 45±15 | 35±14 | 41±13 | 41±13 | 46±14 | 35±15 | 34±9 | 39±5 | 52±8 |
| $SpO_2$ Change (%)[3,4] | | 100±0 | 100±0 | 100±0 | 100±0 | 97±3 | 98±1 | 99±1 | 97±2 | 100±2 | 99±3 | 99±2 | 100+1 | 100±3 | 99±2 | 97±3 | 99±2 |
| Clinical Observations | Responsiveness (#) | 0/6 | 0/6 | 0/6 | 0/4 | 0/6 | 0/6 | 0/6 | 0/4 | 0/3 | 0/3 | 0/3 | 0/2 | 0/3 | 0/3 | 0/3 | 0/2 |
| | Discharges (#) | 0/6 | 0/6 | 0/6 | 0/4 | 0/6 | 0/6 | 0/6 | 0/4 | 0/3 | 0/3 | 0/3 | 0/2 | 0/3 | 0/3 | 0/3 | 0/2 |
| | Skin (#) | 0/6 | 0/6 | 0/6 | 0/4 | 0/6 | 0/6 | 0/6 | 0/4 | 0/3 | 0/3 | 0/3 | 0/2 | 0/3 | 0/3 | 0/3 | 0/2 |
| | Increased Effort in Breathing (#) | 0/6 | 0/6 | 0/6 | 0/4 | **2/6** | **1/6** | **4/6** | **1/4** | **2/3** | **2/3** | **2/3** | **2/2** | **1/3** | 0/3 | **2/3** | **2/2** |
| | Reduced Food Consumption (#) | 0/6 | 0/6 | 0/6 | **1/4** | **5/6** | **5/6** | **5/6** | **3/4** | **3/3** | **2/3** | **3/3** | **2/2** | **3/3** | **3/3** | **3/3** | **2/2** |
| | Soft Stool (#) | **2/6** | 0/6 | 0/6 | 0/4 | **3/6** | **2/6** | **1/6** | **1/4** | **3/3** | **2/3** | **2/3** | **1/2** | **1/3** | 0/3 | 0/3 | 0/2 |
| | Liquid Stool (#) | **1/6** | 0/6 | 0/6 | 0/4 | **1/6** | 0/6 | 0/6 | 0/4 | **1/3** | **1/3** | 0/3 | 0/2 | 0/3 | 0/3 | 0/3 | 0/2 |

n=6 for CoVLP ± adjuvant groups and n=4 for control group at Day 0 pre-challenge and Day 6 post-challenge. n=3 for CoVLP ± adjuvant groups and n=2 at Days 13 and 20 post challenge.
$\delta$: Delta (Change in)
SD: Standard Deviation
#: Occurrence of observations
[1] Pre-Challenge represents data obtained before viral administration on the day of challenge, except for clinical observations for which observations were recorded one day before the challenge.
[2] Clinical signs were observed every day. Day 6 data indicates the incidence observed between Days 0 to 6; Day 13 data indicates the incidence observed between Days 7 to 13; and Day 20 indicates the incidence observed between Days 14 to 20.
[3] Data are presented as mean ± SD.
[4] Changes in temperature, weight and $SpO_2$ are relative to challenge day (indicated as Pre-Challenge).

#### Supplementary Table 4: Post-Challenge Clinical Signs in Rhesus Macaques Infected with SARS-CoV-2 after Two Immunizations with CoVLP Unadjuvanted or Adjuvanted with AS03 or CpG 1018 Adjuvants

| Parameter | | Pre-Challenge[1] | | | | Post-Challenge[2] | | | | | | | | | | | |
|---|---|---|---|---|---|---|---|---|---|---|---|---|---|---|---|---|---|
| | | | | | | Day 6 | | | | Day 13 | | | | Day 20 | | | |
| | | **CoVLP** | **CoVLP + CpG 1018** | **CoVLP + AS03** | **Control** | **CoVLP** | **CoVLP + CpG 1018** | **CoVLP + AS03** | **Control** | **CoVLP** | **CoVLP + CpG 1018** | **CoVLP + AS03** | **Control** | **CoVLP** | **CoVLP + CpG 1018** | **CoVLP + AS03** | **Control** |
| Temp. Change (°C)[3,4] | | 0.0±0.0 | 0.0±0.0 | 0.0±0.0 | 0.0±0.0 | 0.4±0.3 | 0.2±0.5 | 0.4±0.3 | 0.4±1.0 | 0.3±0.5 | 0.1±0.8 | 0.3±0.5 | 0.6±0.4 | 0.1±0.3 | 0.3±0.8 | 0.4±0.6 | 0.0±0.7 |
| $\delta$ Weight (%)[3,4] | | 100±0 | 100±0 | 100±0 | 100±0 | 103±3 | 103±3 | 103±2 | 102±1 | 102±2 | 104±3 | 103±3 | 102±1 | 105±2 | 111±2 | 109±5 | 107±1 |
| Respiratory Rate[3] | | 31±4 | 31±9 | 30±5 | 37±8 | 32±7 | 38±13 | 28±5 | 42±15 | 31±6 | 28±4 | 28±5 | 40±11 | 44±7 | 45±9 | 40±7 | 45±30 |
| $SpO_2$ Change (%)[3,4] | | 100±0 | 100±0 | 100±0 | 100±0 | 100±4 | 98±2 | 98±2 | 97±2 | 100±1 | 97±1 | 101±1 | 97±1 | 100±2 | 99±2 | 101±2 | 101±2 |
| Clinical Observations | Responsiveness (#) | 0/6 | 0/6 | 0/6 | 0/4 | 0/6 | 0/6 | 0/6 | 0/4 | 0/6 | 0/4 | 0/6 | 0/2 | 0/3 | 0/3 | 0/4 | 0/2 |
| | Discharges (#) | 0/6 | 0/6 | 0/6 | 0/4 | 0/6 | 0/6 | 0/6 | 0/4 | 0/6 | 0/4 | 0/6 | 0/2 | 0/3 | 0/3 | 0/4 | 0/2 |
| | Skin (#) | 0/6 | 0/6 | 0/6 | 0/4 | 0/6 | 0/6 | 0/6 | 0/4 | 0/6 | 0/4 | 0/6 | 0/2 | 0/3 | 0/3 | 0/4 | 0/2 |
| | Increased Effort in Breathing (#) | 0/6 | 0/6 | 0/6 | 0/4 | **2/6** | **2/6** | **5/6** | **2/4** | **2/3** | **2/3** | **4/4** | **1/2** | **2/3** | **3/3** | **4/4** | **1/2** |
| | Reduced Food Consumption (#) | **1/6** | **1/6** | **2/6** | **1/4** | **6/6** | **6/6** | **5/6** | **4/4** | **3/3** | **3/3** | **4/4** | **2/2** | **2/3** | **3/3** | **4/4** | **2/2** |
| | Soft Stool (#) | **1/6** | **1/6** | 0/6 | **1/4** | **2/6** | **2/6** | **1/6** | 0/4 | **2/3** | **1/3** | 0/4 | **1/2** | **1/3** | **1/3** | **3/4** | **1/2** |
| | Liquid Stool (#) | **1/6** | 0/6 | 0/6 | 0/4 | 0/6 | 0/6 | 0/6 | 0/4 | 0/3 | 0/3 | **1/4** | 0/2 | 0/3 | 0/3 | 0/4 | 0/2 |

n=6 for CoVLP ± adjuvant groups and n=4 for control group at Day 0 pre-challenge and Day 6 post-challenge. n=3 for CoVLP ± CpG 1018, n=4 for CoVLP + AS03 group and n=2 for Control group at Days 13 and 20 post challenge. Due to logistic reasons, animals were euthanized on Days 20, 21 or 23, results are presented as Day 20.

$\delta$: Delta (Change in).

#: Occurrence of observations.

[1] Pre-Challenge represents data obtained before viral administration on the day of challenge, except for clinical observations for which observations were recorded one day before the challenge.

[2] Clinical signs were observed every day. Day 6 data indicates the incidence observed between Days 0 to 6; Day 13 data indicates the incidence observed between Days 7 to 13; and Day 20 indicates the incidence observed between Days 14 to 20.

[3] Data are presented as mean ± SD.

[4] Changes in temperature, weight and $SpO_2$ are relative to challenge day (indicated as Pre-Challenge).

#### Supplementary Table 5: Peripheral Hematology in Rhesus Macaques Infected with SARS-CoV-2 after One Immunization with CoVLP Vaccine, Unadjuvanted or Adjuvanted with AS03 or CpG 1018 Adjuvants

| Parameter | Pre-challenge[1] | | | | Post-Challenge[2] | | | | | | | | | | | |
|---|---|---|---|---|---|---|---|---|---|---|---|---|---|---|---|---|
| | | | | | Day 6 | | | | Day 13 | | | | Day 20 | | | |
| | CoVLP | CoVLP + CpG 1018 | CoVLP + AS03 | Control | CoVLP | CoVLP + CpG 1018 | CoVLP + AS03 | Control | CoVLP | CoVLP + CpG 1018 | CoVLP + AS03 | Control | CoVLP | CoVLP + CpG 1018 | CoVLP + AS03 | Control |
| WBC ($10^3$/µL) | 8.1±2.4 | 8.0±1.8 | 8.9±2.8 | 7.4±1.2 | 8.3±3.4 | 7.1±1.9 | 8.9±2.9 | 8.2±1.9 | 6.7±1.3 | 8.8±0.8 | 8.9±2.0 | 9.3±1.1 | 6.8±2.3 | 7.4±1.0 | 10.4±2.5 | 8.8±3.5 |
| RBC ($10^6$/µL) | 5.11±0.43 | 5.33±0.39 | 5.38±0.34 | 5.53±0.32 | 5.39±0.19 | 5.34±0.33 | 5.42±0.22 | 5.54±0.26 | 5.53±0.24 | 5.31±0.36 | 5.47±0.21 | 5.28±0.52 | 5.61±0.34 | 5.27±0.29 | 5.44±0.20 | 5.45±0.07 |
| HGB (gm/dL) | 11.7±0.5 | 12.2±0.6 | 12.0±0.6 | 12.7±1.1 | 12.2±0.4 | 12.5±0.6 | 12.3±0.8 | 12.2±0.6 | 12.5±0.4 | 12.5±0.6 | 12.6±0.2 | 11.5±0.1 | 12.7±0.6 | 12.4±0.5 | 12.6±0.2 | 12.1±1.1 |
| Hematocrit (%) | 36.5±1.7 | 38.3±2.0 | 37.9±2.0 | 39.2±3.5 | 37.4±1.2 | 37.7±2.5 | 38.0±2.4 | 37.3±0.9 | 38.1±1.6 | 37.2± 2.2 | 38.4±0.7 | 35.3±1.4 | 39.0±2.2 | 37.1± 2.1 | 38.1±0.5 | 37.1±2.0 |
| MCV (fL) | 71.7±3.3 | 71.9±2.1 | 70.5±2.0 | 70.9±3.9 | 69.5±2.1 | 70.6±1.3 | 70.1±2.8 | 67.5±3.3 | 69.0±0.3 | 70.0±1.5 | 70.3±2.4 | 67.2±3.9 | 69.5±05 | 70.4±1.2 | 70.0±1.6 | 68.1±4.5 |
| MCH (pg) | 22.9±1.1 | 23.0±0.9 | 22.4±0.7 | 23.0±1.3 | 22.7±0.9 | 23.4±0.6 | 22.8±0.8 | 22.1±1.8 | 22.6± 0.4 | 23.7±1.3 | 23.1±0.8 | 21.9±2.4 | 22.7±0.4 | 23.6±1.0 | 23.1±0.8 | 22.1±2.3 |
| MCHC (gm/dL) | 32.0±0.3 | 31.9±0.5 | 31.8±0.3 | 32.5±0.4 | 32.6±0.4 | 33.1±0.6 | 32.5±0.3 | 32.8±1.0 | 32.8±0.4 | 33.7± 1.1 | 32.9±0.0 | 32.6±1.7 | 32.7±0.3 | 33.5±1.0 | 33.1±0.5 | 32.5±1.1 |
| RDW (%) | 14.4±0.8 | 14.1±0.9 | 14.0±0.5 | 13.9±0.7 | 12.6±1.0 | 12.3±1.0 | 12.1±0.2 | 13.9±0.5 | 12.2±1.1 | 12.2±0.7 | 12.0±0.1 | 14.4± 0.1 | 12.3±1.1 | 12.1±0.9 | 11.7±0.2 | 14.3±0.1 |
| Platelet ($10^3$/µL) | 362±54 | 418±68 | 406±60 | 399±45 | 351±136 | 442±95 | 308±95 | 373±138 | 289±105 | 343±52 | 377±171 | 289±134 | 345±103 | 436±60 | 404±121 | 528±25 |
| MPV (fL) | 8.5±0.6 | 8.5±0.8 | 8.4±1.0 | 8.1±0.6 | 11.3±1.4 | 11.3±1.0 | 12.1±0.7 | 11.1±0.2 | 11.9±1.2 | 11.9±1.0 | 11.3±0.5 | 11.3±0.3 | 11.8±0.5 | 11.3±1.1 | 10.9±0.9 | 10.0±0.8 |
| Neut ($10^3$/µL) | 4.3±1.6 | 4.4±1.4 | 5.4±2.0 | 4.4±1.1 | 4.4±2.9 | 3.4±1.3 | 4.4±2.5 | 3.7±1.1 | 3.6±0.9 | 5.3± 1.1 | 5.3±2.0 | 5.3± 1.0 | 3.2±1.3 | 4.1±0.2 | 6.6±1.5 | 4.7±2.1 |
| Lymp ($10^3$/µL) | 2.9±0.9 | 2.9±1.0 | 2.8±1.2 | 2.5±1.0 | 3.4±0.5 | 3.3±1.1 | 4.0±1.2 | 3.9±1.8 | 2.9±0.6 | 3.1±1.0 | 3.2±0.6 | 3.6±0.2 | 3.2±1.0 | 3.0±0.8 | 3.4±1.3 | 3.7±1.5 |
| Mono ($10^3$/µL) | 0.5±0.2 | 0.4±0.1 | 0.4±0.3 | 0.4±0.2 | 0.4±0.3 | 0.4±0.2 | 0.4±0.2 | 0.5±0.2 | 0.2±0.1 | 0.3±0.0 | 0.3±0.1 | 0.3±0.2 | 0.3±0.0 | 0.2±0.0 | 0.4±0.1 | 0.3±0.1 |
| Eos ($10^3$/µL) | 0.2±0.1 | 0.3±0.2 | 0.3±0.3 | 0.2±0.2 | 0.0±0.0 | 0.0±0.0 | 0.0±0.0 | 0.0±0.0 | 0.0±0.0 | 0.0±0.0 | 0.0±0.0 | 0.0±0.0 | 0.0±0.0 | 0.0±0.0 | 0.0±0.0 | 0.1±0.0 |
| Baso ($10^3$/µL) | 0.0±0.1 | 0.0±0.0 | 0.0±0.0 | 0.0±0.0 | 0.0±0.0 | 0.0±0.0 | 0.0±0.0 | 0.0±0.0 | 0.0±0.0 | 0.0±0.0 | 0.0±0.0 | 0.0±0.0 | 0.0±0.0 | 0.0±0.0 | 0.0±0.0 | 0.0±0.0 |

n=12 for CoVLP ± adjuvant groups and n=8 for control group at pre-challenge (Day 7 post immunization). n=5 for CoVLP ± adjuvant groups at Day 6, n= 3 at Days 13 and 21. n=3 for control at Day 6, n=2 at Days 13 and 21.

Data are presented as mean ± SD

SD: Standard Deviation

[1] Pre-challenge analyses were performed at New Iberia Research Center facility.

[2] Animals were transferred to Tulane Primate National Research Center facility for SARS-CoV-2 infection, where further analyses were performed.

**Supplementary Table 6: Peripheral Hematology in Rhesus Macaques Infected with SARS-CoV-2 after Two Immunizations with CoVLP Vaccine, Unadjuvanted or Adjuvanted with AS03 or CpG 1018 Adjuvants**

| Parameter | Pre-challenge[1] | | | | Post-Challenge[2] | | | | | | | | | | | |
|---|---|---|---|---|---|---|---|---|---|---|---|---|---|---|---|---|
| | | | | | Day 6 | | | | Day 13 | | | | Day 20 | | | |
| | CoVLP | CoVLP + CpG 1018 | CoVLP + AS03 | Control | CoVLP | CoVLP + CpG 1018 | CoVLP + AS03 | Control | CoVLP | CoVLP + CpG 1018 | CoVLP + AS03 | Control | CoVLP | CoVLP + CpG 1018 | CoVLP + AS03 | Control |
| WBC ($10^3$/μL) | 7.0±1.8 | 7.0±2.5 | 7.2±1.9 | 8.3±1.8 | 9.8±3.8 | 8.3±2.2 | 9.6±2.3 | 8.4±3.0 | 11.3±4.2 | 9.3±3.0 | 7.9±2.5 | 9.1±2.9 | 9.1±0.7 | 8.4±1.3 | 10.4±2.6 | 9.8±0.3 |
| RBC ($10^6$/μL) | 5.20±0.38 | 5.28±0.49 | 5.51±0.42 | 5.67±0.29 | 5.07±0.45 | 5.35±0.40 | 5.66±0.43 | 5.44±0.13 | 5.41±0.36 | 5.51±0.54 | 5.76±0.50 | 5.59±0.17 | 5.70±0.16 | 5.98±0.48 | 5.80±0.54 | 5.88±0.08 |
| HGB (gm/dL) | 11.9±0.4 | 12.2±0.8 | 12.3±1.0 | 13.5±0.7 | 11.8±0.7 | 12.5±0.9 | 12.7±0.8 | 13.3±0.7 | 12.4±0.4 | 12.9±0.8 | 13.1±0.8 | 13.5±0.1 | 13.0±0.5 | 14.2±0.6 | 13.3±0.9 | 14.3±0.2 |
| Hematocrit (%) | 37.4±1.5 | 38.0±2.5 | 38.5±2.8 | 41.6±1.5 | 35.8±2.0 | 37.7±2.5 | 38.9±2.5 | 39.0±1.4 | 36.8±1.7 | 38.8±2.8 | 39.5±2.7 | 39.6±0.1 | 38.8±1.2 | 42.2±2.7 | 39.7±3.0 | 41.9±1.0 |
| MCV (fL) | 72.2±3.2 | 72.1±2.9 | 69.9±2.1 | 73.6±1.9 | 70.7±3.4 | 70.7±1.7 | 68.8±1.1 | 71.6±1.8 | 68.2±1.7 | 70.6±2.1 | 68.6±1.5 | 70.9±1.9 | 68.0±1.7 | 70.6±1.7 | 68.6±1.5 | 71.4±2.6 |
| MCH (pg) | 23.0±0.9 | 23.2±1.0 | 22.4±0.9 | 23.8±0.9 | 23.3±1.1 | 23.5±1.0 | 22.5±0.9 | 24.4±0.8 | 22.9±1.0 | 23.5±0.9 | 22.7±0.7 | 24.1±0.6 | 22.8±0.9 | 23.7±1.1 | 23.1±1.0 | 24.3±0.6 |
| MCHC (gm/dL) | 31.8±0.4 | 32.2±0.3 | 32.0±0.3 | 32.4±0.6 | 32.9±0.6 | 33.2±0.7 | 32.8±1.0 | 34.0±0.6 | 33.6±0.6 | 33.3±0.2 | 33.2±0.4 | 34.0±0.1 | 33.5±0.5 | 33.6±0.8 | 33.6±0.8 | 34.0±0.3 |
| RDW (%) | 14.3±0.9 | 14.1±0.5 | 14.1±0.6 | 13.8±0.3 | 12.2±0.4 | 12.1±0.4 | 12.3±0.7 | 12.3±0.9 | 12.2±0.2 | 12.3±0.8 | 12.4±0.5 | 12.4±0.6 | 12.1±0.1 | 12.1±0.8 | 12.3±0.4 | 12.3±0.6 |
| Platelet ($10^3$/μL) | 364±84 | 391±76 | 411±34 | 453±42 | 342±53 | 367±60 | 308±109 | 442±111 | 312±94 | 393±100 | 294±91 | 421±70 | 405±54 | 466±75 | 434±54 | 556±58 |
| MPV (fL) | 8.4±0.6 | 8.7±0.8 | 8.6±1.0 | 8.1±0.5 | 11.6±0.5 | 11.6±1.0 | 12.0±1.1 | 11.5±0.5 | 11.3±0.6 | 11.2±1.0 | 11.5±0.7 | 11.2±0.4 | 10.7±0.7 | 10.8±0.6 | 10.3±0.6 | 11.3±0.5 |
| Neut ($10^3$/μL) | 3.6±1.0 | 3.8±2.4 | 4.2±1.6 | 5.1±1.8 | 4.8±2.3 | 4.8±1.6 | 5.5±2.1 | 4.9±2.4 | 5.9±1.5 | 6.8±2.3 | 4.1±1.8 | 6.5±2.0 | 5.4±1.6 | 5.7±1.7 | 6.8±3.0 | 6.5±0.5 |
| Lymp ($10^3$/μL) | 2.7±0.9 | 2.5±0.6 | 2.3±0.7 | 2.6±0.7 | 4.1±1.6 | 2.9±0.8 | 3.6±1.1 | 2.8±0.6 | 4.8±2.5 | 2.2±0.8 | 3.5±1.9 | 2.4±1.0 | 3.2±0.7 | 2.3±0.5 | 3.3±1.1 | 3.0±0.7 |
| Mono ($10^3$/μL) | 0.4±0.2 | 0.4±0.2 | 0.3±0.1 | 0.4±0.2 | 0.8±0.3 | 0.6±0.1 | 0.5±0.3 | 0.6±0.3 | 0.4±0.1 | 0.3±0.1 | 0.2±0.1 | 0.1±0.1 | 0.4±0.1 | 0.4±0.2 | 0.3±0.1 | 0.3±0.0 |
| Eos ($10^3$/μL) | 0.3±0.1 | 0.2±0.1 | 0.4±0.2 | 0.2±0.1 | 0.0±0.0 | 0.0±0.0 | 0.0±0.0 | 0.0±0.0 | 0.1±0.1 | 0.0±0.0 | 0.1±0.1 | 0.0±0.0 | 0.1±0.1 | 0.0±0.0 | 0.0±0.0 | 0.1±0.0 |
| Baso ($10^3$/μL) | 0.0±0.1 | 0.0±0.0 | 0.0±0.0 | 0.0±0.0 | 0.0±0.0 | 0.0±0.0 | 0.0±0.0 | 0.0±0.0 | 0.0±0.0 | 0.0±0.0 | 0.0±0.0 | 0.0±0.0 | 0.0±0.0 | 0.0±0.0 | 0.0±0.0 | 0.0±0.0 |

n=6 for CoVLP ± adjuvant groups and n=4 for control group at pre-challenge (Day 7 after the second immunization) and Day 6 post-challenge. n=3 for CoVLP ± CpG 1018 groups, n=4 for CoVLP + AS03 group and n=2 for control group at Days 13 and 20 post infection. Due to logistic reasons, animals were euthanized on Days 20, 21 or 23, results are presented as Day 20.

Data are presented as mean ± SD

[1] Pre-challenge analyses were performed at New Iberia Research Center facility.

| Parameter | Pre-challenge[1] | | | | Post-Challenge[2] | | | | | | | | | | | |
|---|---|---|---|---|---|---|---|---|---|---|---|---|---|---|---|---|
| | | | | | Day 6 | | | | Day 13 | | | | Day 20 | | | |
| | CoVLP | CoVLP + CpG 1018 | CoVLP + AS03 | Control | CoVLP | CoVLP + CpG 1018 | CoVLP + AS03 | Control | CoVLP | CoVLP + CpG 1018 | CoVLP + AS03 | Control | CoVLP | CoVLP + CpG 1018 | CoVLP + AS03 | Control |

[2] Animals were transferred to Tulane Primate National Research Center facility for SARS-CoV-2 infection, where further analyses were performed.

#### ***Appendix***

Date: ________________ **Cage-Side Assessment** Animal ID: ________________

| Parameter | Description | Score |
|---|---|---|
| Responsiveness | Normal - bright, alert, responsive | 0 |
| | Mildly affected - slightly depressed, acts disinterested with personnel in room, lies down in cage but gets up when approached | 1 |
| | Moderately affected/obtunded - non-responsive, very disinterested in personnel, hunched or lying down, will get up when stimulated | 2 |
| | Severely affected/comatose - lying down completely unresponsive to stimuli | 3 |
| Discharges | Normal | 0 |
| | Mild nasal/ocular | 1 |
| | Severe nasal/ocular | 3 |
| Skin | Normal | 0 |
| | Mild dermatitis | 1 |
| | Severe dermatitis | 3 |
| Respiratory | Normal - no apparent changes in breathing, (est 16-54 BPM), no cough | 0 |
| | Mild - slightly increased effort breathing, (est 55-66 BPM), and/or mild cough | 1 |
| | Moderate - obvious difficulty breathing, (est 67-80 BPM), and/or moderate cough | 2 |
| | Severe - open mouth breathing, abdominal breathing, (est >80 BPM), cyanosis, and/or severe cough | 3 |
| Food consumption | <25% of food remaining | 0 |
| | ≤25%-50% of food remaining | 1 |
| | >50% of food remaining | 2 |
| Fecal consistency | Normal | 0 |
| | Soft | 1 |
| | Fluid | 2 |

| |
|---|
| Total |
| Notes *(any observed sneezing, vomit, conjunctival erythema, or other abnormalities*) |

Date: ________________ **Physical Examination Under Anesthesia** Animal ID: _________________

| **Parameter** | **Description** | **Score** |
|---|---|---|
| Rectal temperature *(taken immediately after anesthesia induction)* | Normal (100.0-102.4F) | 0 |
| | Mild hypothermia (98.0-99.9F) | 1 |
| | Moderate hypothermia (96.0-97.9F) | 2 |
| | Severe hypothermia (≤96.0) | 3 |
| | Mild hyperthermia (102.5-103.4F) | 1 |
| | Moderate hyperthermia (103.5-104.4F) | 2 |
| | Severe hyperthermia (>104.4F) | 3 |
| Respiratory rate | Normal (16-54 BPM) | 0 |
| | Mild tachypnea (55-66 BPM) | 1 |
| | Moderate tachypnea (67-80 BPM) | 2 |
| | Severe tachypnea (>80 BPM) | 3 |
| Respiratory character | Normal | 0 |
| | Mild dyspnea | 1 |
| | Severe dyspnea | 3 |
| Auscultation | Normal | 0 |
| | Mild (occasional crackles/rales or wheezes) | 1 |
| | Severe (continuous crackles/rales or wheezes) | 3 |
| $SpO_2$ | Normal (96-100%) | 0 |
| | Mildly decreased (92-95) | 1 |
| | Moderately decreased (80-91) | 2 |
| | Severely decreased (<80%) | 3 |
| | Normal (0-4.9% loss) | 0 |

| | | |
|---|---|---|
| Body weight | Mild (5-10.9% loss) | 1 |
| | Moderate (11-24.9% loss) | 2 |
| | Severe (≥25% loss) | 3 |
| Hydration | Normal (normal skin turgor, moist mucous membranes) | 0 |
| | Mild dehydration (5-10%) | 1 |
| | Severe dehydration (>10%) | 3 |
| **Notes** | | **Total** |
